# Cell layer specific roles for hormones in root development: Gibberellins suppress infection thread progression, promote nodule and lateral root development in the endodermis and interact with auxin and cytokinin

**DOI:** 10.1101/2023.09.04.555035

**Authors:** Karen Velandia, Alejandro Correa-Lozano, Peter M. McGuiness, James B. Reid, Eloise Foo

## Abstract

1. Gibberellins have a profound influence on the formation of lateral root organs. However, the precise role this hormone plays in the cell-specific events during lateral root formation, rhizobial infection and nodule organogenesis, including interactions with auxin and cytokinin, is not clear.
2. We performed epidermal- and endodermal-specific complementation of the severely gibberellin-deficient *na* pea (*Pisum sativum*) mutant with *Agrobacterium rhizogenes*. Gibberellin mutants were used to examine the spatial expression pattern of cytokinin (*TCSn*) and auxin (*DR5*) responsive promoters and hormone levels.
3. We found that gibberellins produced in the endodermis promote lateral root and nodule organogenesis and can induce a mobile signal(s) that suppresses rhizobial infection. In contrast, epidermal-derived gibberellins suppress infection but have little influence on root or nodule development. Gibberellins suppress the cytokinin-responsive *TCSn* promoter in the cortex and are required for normal auxin activation during nodule primordia formation.
4. Our findings indicate that gibberellins regulate the checkpoints between infection thread penetration of the cortex and invasion of nodule primordial cells and promotes the subsequent progression of nodule development. It appears that gibberellins limit the progression and branching of infection threads in the cortex by restricting cytokinin response and activate auxin response to promote nodule primordia development.

## Introduction

The formation of *de novo* root organs, such as lateral roots and nodules, involves precise spatial and temporal changes in cell division and differentiation. Until recently, much of our knowledge of lateral root development was limited to studies in *Arabidopsis*, which indicated an essential role for the pericycle in lateral root emergence (Chiatante *et al*., 2018). We now know that brassicas are likely an exception and that most seed plants derive lateral roots from a much broader group of cells, including the pericycle, cortex and endodermis (Xiao *et al*., 2019). Selected species in the fabid clade also form unique lateral organs, nodules, to host nitrogen-fixing bacteria. Nodules are formed from the inner cell layers of the root in legumes forming indeterminate nodules and appear to be derived in part from the lateral root program (Schiessl *et al*., 2019; Soyano *et al*., 2019; Shrestha *et al*., 2021; Soyano *et al*., 2021). Infection occurs in the epidermis and requires precise spatial and temporal co-ordination to enable colonisation of developing nodules in the inner cortex. Plant hormones play key spatial and temporal roles in driving the development of roots and nodules (Chiatante *et al*., 2018; Velandia *et al*., 2022). In this study using the model legume pea, we demonstrate a powerful role of gibberellins (GA) derived from the endodermis in promoting lateral root and nodule development and reveal complex interaction between gibberellins, auxin and cytokinins.

Gibberellins are a group of diterpenoid carboxylic acids, with only a small number of them being bioactive (e.g. GA_1_, GA_3_ and GA_4_) in any one species (Hedden, 2016). The biosynthesis and catabolism pathways are tightly controlled by a series of enzymes with complex and sometimes tissue-specific expression patterns (Davidson *et al*., 2003; Hedden, 2020). In soybean (*Glycine max*) and *Medicago truncatula*, gene expression and promoter::*GUS* studies have identified genes coding for specific gibberellin biosynthesis (GA3-oxidation and GA20-oxidation) and catabolism (GA2-oxidation) enzymes whose expression is specifically regulated in response to rhizobial inoculation and co-occurs in cells undergoing infection and/or nodule development (Kim *et al*., 2019; Chu *et al*., 2021; Gao *et al*., 2023). In *Medicago* the expression of *DELLA* genes, that encode transcriptional repressors of gibberellin response that are targeted for degradation by gibberellin perception, are also elevated in response to infection and in nodule tissue (Fonouni-Farde *et al*., 2016; Jin *et al*., 2016). Although it is not possible to predict the actual level of bioactive gibberellin from these studies, as gibberellin biosynthesis and catabolism gene expression are under strong feedback control (Reid *et al*., 2002; Hedden & Sponsel, 2015), the use of a gibberellin responsive biosensor in soybean indicates elevated gibberellin response occurs during infection and in actively dividing and expanding cells of nodules (Chu *et al*., 2021) .

Careful analysis of the phenotypes of gibberellin-treated plants and lines with altered gibberellin biosynthesis or signalling reveal a complex role for gibberellins during nodulation. Gibberellin suppresses rhizobial infection (Maekawa *et al*., 2009; Fonouni-Farde *et al*., 2016; Jin *et al*., 2016; McAdam *et al*., 2018; Kim *et al*., 2019; Gao *et al*., 2023). In contrast, there is significant evidence that gibberellins play a positive role in nodule organogenesis and development. Nodule number, size and in some cases function, is negatively impacted in plants with disrupted gibberellin biosynthesis in pea, soybean and *Medicago truncatula* (Ferguson *et al*., 2005; McAdam *et al*., 2018; Kim *et al*., 2019; Chu *et al*., 2021; McGuiness *et al*., 2021; Gao *et al*., 2023). Furthermore, *Psdella* mutants develop less nodules, likely due to reduced number of infections, but these nodules display normal size and function (McAdam *et al*., 2018). In *Medicago* a somewhat different result was found, with a range of nodule types observed in mutants missing all *DELLA* genes, ranging from normal to smaller than wild type (WT) (Jin *et al*., 2016) .

Clearly, in studies examining the role of gibberellins in rhizobial-induced nodules it is essential to consider both the effects on infection and nodule organogenesis, as gibberellin-induced reduction in infection could indirectly lead to a reduction in nodule number. One way to address this is to examine nodule-like structures that can be induced in the absence of infection by rhizobia. Indeed, exogenous application of gibberellin induces the division of pericycle cells to form nodule-like structures in some legumes and this appears to require Nodule Inception (NIN) (Kawaguchi *et al*., 1996; Akamatsu *et al*., 2021) . In contrast, in *Medicago* the expression of a *della* dominant active protein induces spontaneous nodule-like structures and gibberellin application or loss of function *della* mutants inhibit spontaneous nodule formation in *ccamk* and cytokinin receptor gain of function mutants (Maekawa *et al*., 2009; Fonouni-Farde *et al*., 2016; Jin *et al*., 2016) . It is difficult to compare these nodule-like structures across studies, as not all species respond in the same way and detailed analysis of the cellular structure of these growths is not always reported. More detailed studies, using cell-specific approaches, are required to clarify the dual role of gibberellins in infection and nodule development.

Like gibberellin, cytokinin plays distinct roles in the different cell layers, inhibiting infection but promoting nodule organogenesis and development, while auxin appears to promote both stages of nodulation (Velandia *et al*., 2022). Studies have indicated a possible feedback mechanism where cytokinin produced in the epidermis may suppress infection but enhance nodule development in the cortex (Jardinaud *et al*., 2016; Reid *et al*., 2016; Reid *et al*., 2017; Jarzyniak *et al*., 2021), while cytokinin produced in the cortex may suppress infection in the epidermis (Reid *et al*., 2016; Miri *et al*., 2019). Although it is not clear if cytokinin itself is moving, there is evidence that cytokinin may be transported in *Medicago* from the epidermis to the cortex to promote nodule organogenesis (Jarzyniak *et al*., 2021). Additionally, cytokinin seems to be required for the subsequent auxin accumulation during nodule organogenesis (Plet *et al*., 2011; Rightmyer & Long, 2011). In pea, there is relatively little knowledge about the role of cytokinin in nodulation. High cytokinin levels and cytokinin-responsive gene expression are localised in developing pea nodules (Dolgikh *et al*., 2020). Interestingly, application of the synthetic cytokinin 6-Benzylaminopurine (BAP) can suppress nodule number (Lorteau *et al*., 2001; Fei & Vessey, 2003) and induce highly branched infection threads (Lorteau *et al*., 2001). Previous studies in pea using double mutants have found gibberellins promote nodule initiation via suppression of ethylene biosynthesis, but that these two hormones act relatively independently to suppress infection and to promote nodule maturation (Ferguson *et al*., 2011; McAdam *et al*., 2018). Our understanding of how gibberellins interact with auxin and/or cytokinins during nodulation is relatively limited (Weston *et al*., 2009; Fonouni-Farde *et al*., 2017; Fonouni-Farde *et al*., 2018). One powerful way to examine the interaction between gibberellins, auxin and cytokinins is the use of hormone biosensors, as previous studies have revealed tight spatial control of auxin and cytokinin response during infection and nodule development (Ng & Mathesius, 2018; Nadzieja *et al*., 2019).

In this study, we examined cell-specific effects of gibberellin synthesis on infection, nodule and lateral root development and explored the interaction between auxin, cytokinin and gibberellin using cytokinin- and auxin-responsive biosensors, hormone application and hormone quantification. Mutants disrupted in key gibberellin biosynthesis enzymes including NA (ent-kaurenoic acid oxidase; Davidson *et al*., 2003) and SLN (2-oxidase; Lester *et al*., 1999; Martin *et al*., 1999) were employed. As gibberellin precursors can be mobile (Binenbaum *et al*., 2018) and gibberellin may influence other mobile elements (Akamatsu *et al*., 2021) the use of cell-layer specific hairy root transformation enabled us to examine local communication between root cell layers and more long-distance communication between roots and shoots.

## Materials and Methods

### Plant material and rhizobial growth

The *Pisum sativum* lines used in this study were the GA deficient mutant *na-1* (Davidson *et al*., 2003), derived from WT line WL1769 and crossed onto the cv. Torsdag background. Wild type and gibberellin deficient *na/na* tagged with the *DR5::GUS* reporter were produced from a cross between *DR5::GUS* on the Cameor background (DeMason & Polowick, 2009) and *na/na* on the cv. Torsdag background. *DR5::GUS* tagged segregants that were homozygous LE/LE were selected in the 3^rd^ and 4^th^ generation. The supernodulating *Psnark* (P88) mutant (Sagan & Duc, 1996) and the corresponding WT Frisson and GA-deficient *sln* mutant and corresponding WT SLN (Reid *et al*., 1992) were also used. Soybean (*Glycine max)* seed was purchased from Range View Seeds (Winnaleah, Tasmania, Australia). Long Ashton nutrients were applied weekly with modification stated for each experiment (Hewitt, 1966). *Rhizobium leguminosarum* symbiovar *viciae* strain RLV3841 carrying pXLGD4 plasmid containing a lacZ reporter and strain RLV248 containing the plasmid Phc60 for GFP constitutive expression (RLV248G) (Cheng & Walker, 1998) were used. RLV248GFP was provided by Maria Soto (University of Granada).

### Construct assembly

Specific promoters and genes where inserted in the Pcambia_CR1 (PCR1) vector (Sevin-Pujol *et al*., 2017) using Golden Gate system. PCR fragments for promoter regions and genes where generated using primers containing the BsaI recognition site and matching overhangs (Table. S2), following the instructions of Golden Gate assembly kit (New England Biolabs). The *PsNA* coding region was obtained from pea root cDNA.

A pea epidermal expression promoter was identified for this study, based on the promoter region from the pea orthologue (Psat0s301g0160) of *AtEXPA7* and *MtEXPA7* genes (Kim *et al*., 2006). A fragment 410bp upstream of the ATG start codon was selected. The *AtCASP1* endodermal promoter (Roppolo *et al*., 2011), was provided by Sandra Bensmihen (Sevin-Pujol *et al*., 2017). The two component signalling (*TCSn*) cytokinin responsive promoter was supplied by Bruno Muller (Zürcher *et al*., 2013). For the *NA* complementation experiments, plasmids containing Promoter::β-glucuronidase (*GUS*) were used as a control.

### Hairy root transformation

Pea roots were transformed with *Agrobacterium rhizogenes* (ARqua1) carrying the corresponding construct following a previously described method (Clemow *et al*., 2011) with some modifications (Methods S1).

For harvesting, individual roots generated from the callus of 10-20 plants were checked for transformation using the DsRED fluorescent marker. For hormone analysis, a complete transformed root was stored in 80% MeOH at -20°C. For RNA extraction, a complete transformed root was stored at -80°C. Roots were stained for GUS before fixation in paraformaldehyde as described in Methods S3. The remaining roots were fixed in 1% paraformaldehyde and stored at 4°C for scoring.

Roots were examined using a ZEISS Axioscope5 fluorescence microscope. The total number of infection threads (IT), IT connected to a nodule primordium, branched IT, developing nodules and mature nodules were counted. Cell length is the average of 25 independent measures and root width is the average of 5 independent measures in the mature zone of the root. Number of cells was calculated according to the root length. For lateral root length, the average of the longest 6 lateral roots was calculated. For *TCSn::GUS* transformation experiments, only the infection zone of the roots (tip to 4cm) was scored for nodulation structures, including the presence of GUS staining associated with each structure.

### Gene expression and RT-PCR

Root RNA was extracted from 5 biological replicates, using ISOLATE II RNA mini kit (Bioline). cDNA was synthetized from 1μg of RNA using SensiFAST cDNA Synthesis kit (Bioline). RT PCR was carried out in a Rotor-gene Q2 PLEX (QIAGEN), using SensiMix SYBR master mix (Bioline), in duplicate for each biological replicate. TFIIa and actin genes were used to calculate the relative expression. Primers for *NA* amplify only the WT version of the gene (Table S2).

### Dynamics of infection and hormone levels

Plants were grown in cabinets, for 6-13 days in sterile conditions and then inoculated with tagged rhizobia as described by McAdam *et al*. (2018). For *na* and *Psnark* experiments, the interzone (2cm of tissue taken 1cm back from root tip) of the root was harvested in 80% MeOH for hormone extraction (Fig. S4 only) and the whole root was stored in 4% paraformaldehyde for scoring on specific days post inoculation (dpi). For *sln* experiments, plants were harvested 3 weeks post inoculation and scored for nodulation structures in the top 40mm and the 40-140mm tap root sections.

### Plant hormone application

To monitor the induction of nodule-like structures in the absence of rhizobia in soybean, seeds were sterilised as outlined above and planted in cabinets in vermiculite in sterile conditions. Plants were grown in 25°C day, 15°C night and 18h photoperiod. To test the effect of GA_3_ or Paclobutrazol (PAC) in pseudonodule formation induced by 6-Benzylaminopurine (BAP), soybean seeds were treated with 20μg and 40μg of GA_3_ or 30μg and 100μgPAC. 75mL of 20μM BAP in nutrient solution was applied weekly by root drench. Plants received weekly nutrients without nitrogen (McAdam *et al*., 2018) and were harvested at day 28. To induce nodule-like structures in pea, WT and *na* plants were grown on slants as described by Blake *et al*. (2016). 9 day old plants received a drench of water with 100uM Naphthylphthalamic acid (NAP) dissolved in MeOH or same volume of MeOH in water as a control and harvested 12 days later. The number of pseudonodules was recorded and expressed per g DW root.

To test the effect of BAP in pea WT, plants were grown and inoculated with lacZ tagged rhizobia. Plants were root drenched weekly with 75mL of 80μM BAP or water and harvested 3 weeks after planting. Plants were scored for infection threads, branched IT and nodules.

### Hormone analysis

Auxin and cytokinin extraction and quantification was carried out as outlined in Bound *et al*. (2022) and detailed in Methods S2, with stable isotope-labelled internal standards were added to each sample. For gibberellin quantification samples were extracted as outlined in Ford *et al*. (2018), with stable isotope-labelled internal standards added to each sample. Samples were then derivatized as described Li *et al*. (2017) except that samples were incubated in 50μL of 200 mM EDC (N-(3-dimethylaminopropyl)-N9-ethylcarbodiimide hydrochloride; Sigma-Aldrich) in ethanol over-night at 40°C, then dried and resuspended in 30μL Milli-Q water. Samples were then analysed using a Waters Acquity H-Class UPLC instrument coupled to a Waters Xevo triple quadrupole mass spectrometer. For all samples, peak areas for endogenous and labelled hormones were compared and combined with the FW of samples to calculate ng/g FW.

## Results

### Cell-specific rescue of gibberellin-deficient mutant

Previous studies have shown that the severely gibberellin-deficient mutant *na* has a dramatic increase in number of infection threads, many of which form highly branched structures in the cortex that are not connected to nodules (McAdam *et al*., 2018). Even though there is a dramatic increase in infection, *na* mutants develop few nodules and the nodules that do develop are not mature and have limited function compared to wild type nodules (Ferguson *et al*., 2005; Ferguson *et al*., 2011; McAdam *et al*., 2018; McGuiness *et al*., 2021). The *na* mutants also have severely dwarfed shoots and roots compared to wildtype (Yaxley *et al*., 2001; Foo *et al*., 2013) . To examine the influence of gibberellin produced in specific layers of the root on these phenotypes, we employed two different cell-layer specific promoters to drive the expression of the wild-type *NA* gene in wild type and *na* mutant roots. For epidermal expression we employed the *PsEXPA* promoter that shares significant homology with Medicago *EXPA7* (Kim *et al*., 2006; Vernie *et al*., 2015), and displays epidermal specific expression in pea roots when linked to *GUS* gene (Fig. S1a-b). We also employed the *AtCASP* promoter (Roppolo *et al*., 2011; Sevin-Pujol *et al*., 2017) that when linked to *GUS* displayed strong and specific endodermal expression in pea roots (Fig. S2a-b).

Epidermal expression of wild type *NA* resulted in a significant increase in the level of *NA* transcript in wild type and *na* transformed roots (Fig S1c). *NA* expression in the epidermis does not appear to be a rate limiting step for gibberellin biosynthesis in wild type roots, as there was no significant effect of *EXPA::NA* expression on rhizobial infection or nodule number compared to *EXPA::GUS* control wild type roots (Fig.1). *EXPA::GUS na* control roots displayed previously reported phenotypes of dramatically increased infection, presence of branched infection threads in the cortex and reduced number of mature nodules compared to wild type roots (Fig. 1). *EXPA::NA* expression in *na* mutant roots resulted in a significant approximately 60% reduction in the number of infection threads compared to *na EXPA::GUS* control roots, although infection was still somewhat elevated compared to wild type (Fig. 1a-c, j). There was no significant effect on other stages of infection or nodulation in *na EXPA::NA* roots compared to *na EXPA::GUS* (Fig.1), with *na EXPA::NA* roots developing small nodules containing a reduced level of rhizobia compared to wild type roots (Fig. 1g-i).

**Fig. 1.**
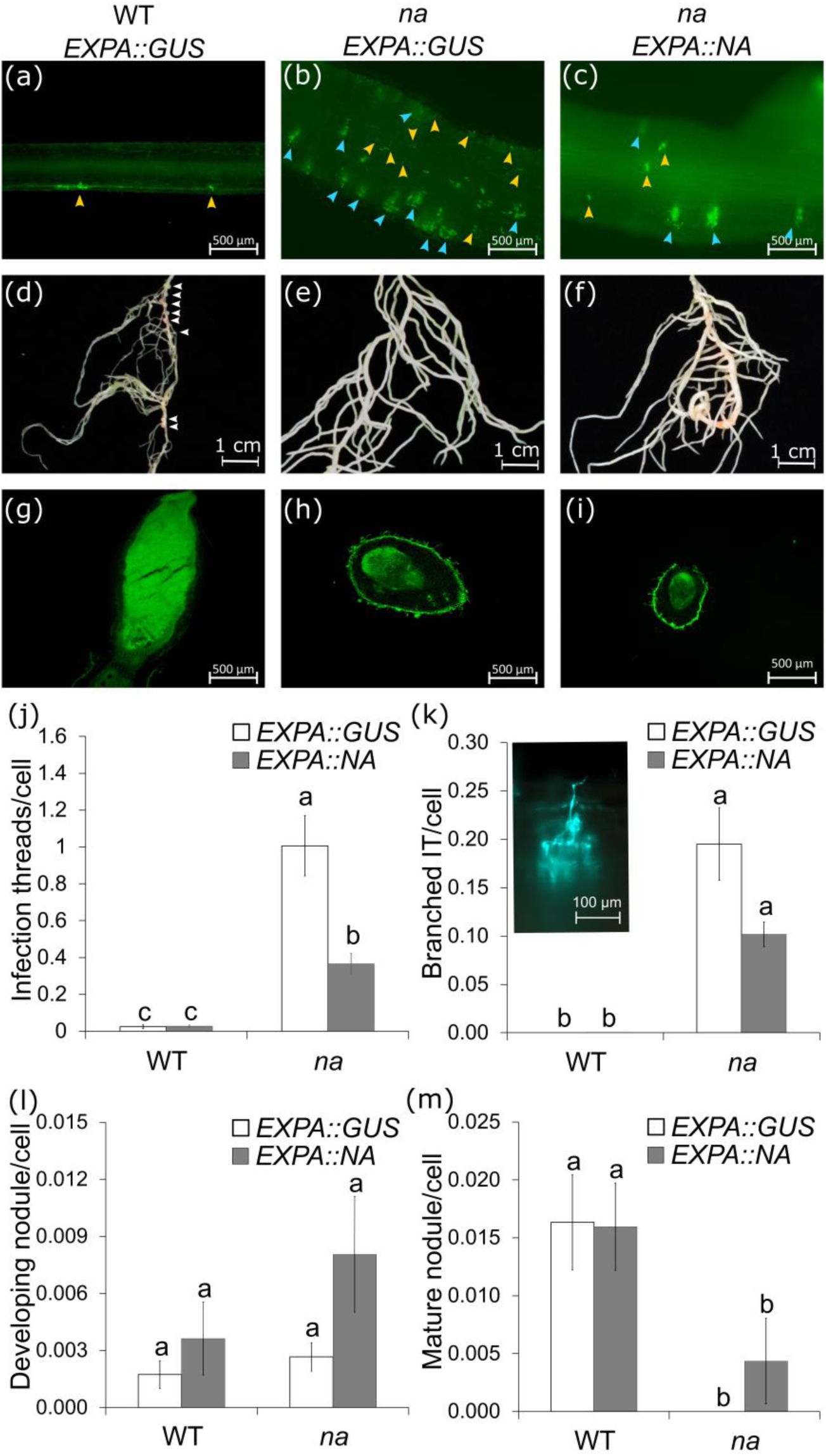
Epidermal expression of the *NA* gene in WT and *na* mutant lines. (a-c) Representative picture of normal (yellow arrowheads) and branched (blue arrowheads) infection threads (rhizobia are shown in green fluorescence) in (a) WT *EXPA::GUS*, (b) *na EXPA::GUS* and (c) *na EXPA::NA.* Note that the images of WT *EXPA::NA* are not shown because the phenotype is not different to WT *EXPA::GUS*. (d-f) Representative picture of transgenic roots, white arrowheads point to mature nodules. (g-i) 40μm sections of a typical nodule. Scoring of (j) infection threads not connected to a nodule primordium, (k) representative photo and scoring of branched infection threads, (l) developing nodules and (m) mature nodules. Values are mean and bars represent the standard error, grouping letters were obtained using one way ANOVA with Tukey *post hoc* test (j,l) and Kruskal-Wallis nonparametric test (k,m). Values with different letters are significantly different (P<0.05), (n = 7-12).

Endodermal expression of wildtype *NA* also resulted in a significant increase in the level of *NA* transcript in wild type and *na* transformed roots (Fig. S2a). Similar to the epidermal experiment, endodermal expression of *NA* in wild type roots did not result in any significant effect on rhizobial infection or nodule number compared to *CASP::GUS* wild type control roots (Fig. 2). Control *na CASP::GUS* roots displayed increased infection, the presence of branched infection threads and a complete absence of mature nodules compared to wild type roots (Fig. 2). *CASP::NA* expression in *na* mutant roots resulted in a significant reduction of 70% in the number of infections in the epidermis compared to *na* control roots (Fig. 2j). *CASP::NA* expression in *na* mutant roots also significantly decreased the number of branched infection threads in the cortex compared to *na CASP::GUS* control roots (Fig. 2k). Strikingly, *na CASP::NA* roots developed many more nodules than *na CASP::GUS* control roots, many of which developed to a mature size (Fig. 2i, l, m). Indeed, *na CASP::NA* roots developed a similar number of mature nodules to wild type roots and with a similar size and amount of rhizobial colonisation to wild type nodules (Fig. 2 g, i, m).

**Fig. 2.**
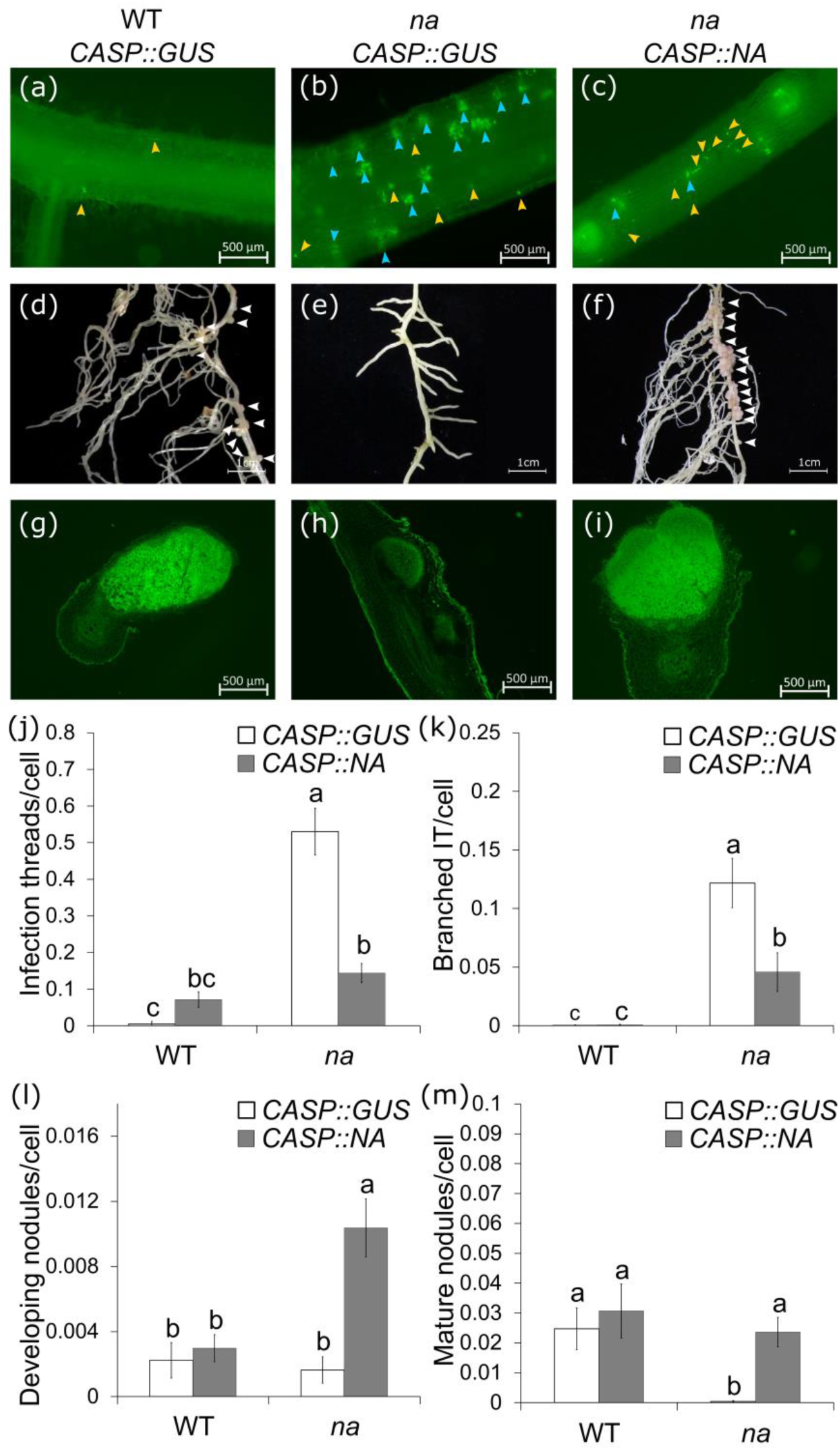
Endodermal expression of the *NA* gene in WT and *na* mutant lines. (a-c) Representative picture of normal (yellow arrowheads) and branched (blue arrowheads) infection threads (rhizobia are shown in green fluorescence) in (a) WT *CASP::GUS*, (b) *na CASP::GUS* and (c) *na CASP::NA.* (d-f) Representative picture of transgenic roots, white arrowheads point to mature nodules. (g-i) 40μm sections of a typical nodule. Scoring of (j) infection threads not connected to a nodule primordium, (k) branched infection threads, (l) developing nodules and (m) mature nodules. Values are mean and bars represent the standard error. Grouping letters were obtained using one way ANOVA with Tukey *post hoc* test (j,k,l) and Kruskal-Wallis non parametric test (m). Values with different letters are significantly different (P<0.05), (n = 5-7). This experiment was repeated three times with similar results.

Root and shoot development were also monitored in the epidermal and endodermal *NA* complementation experiments outlined above (Fig. 3, S1d and S2d). As seen for nodulation, ectopic expression of *NA* in the epidermis or endodermis did not result in any change in root or shoot characteristics of wild type plants. Mutant *na* plants transformed with the control constructs displayed a significant reduction in shoot fresh weight (Fig. S1d, S2d), root cell length, and the number and length of lateral roots and a significant increase in root width compared to wild type (Fig. 3). *EXPA::NA* expression in *na* mutant roots had no significant impact on any of these root or shoot characteristics. In contrast, endodermal *CASP::NA* expression in *na* mutant roots resulted in a significant increase in cell length and lateral root length, and a small, but not significant increase in the number of lateral roots and decrease in root width compared to *CASP::GUS na* roots (Fig.3). Indeed, all of these root characteristics were not significantly different to levels seen in at least one of the wild type treatments. Transformed *EXPA::NA* and *CASP::NA na* roots did not influence the development of untransformed shoots and roots on the same *na* mutant plant, as these untransformed shoots and roots still displayed dwarf shoot and root phenotypes (Fig. S1d, e, S2d, e). This indicates ectopic expression of *NA* in *na* mutants did not induce the long-distance root-to-root or root-to-shoot transport of bioactive gibberellins or other active mobile elements.

**Fig. 3.**
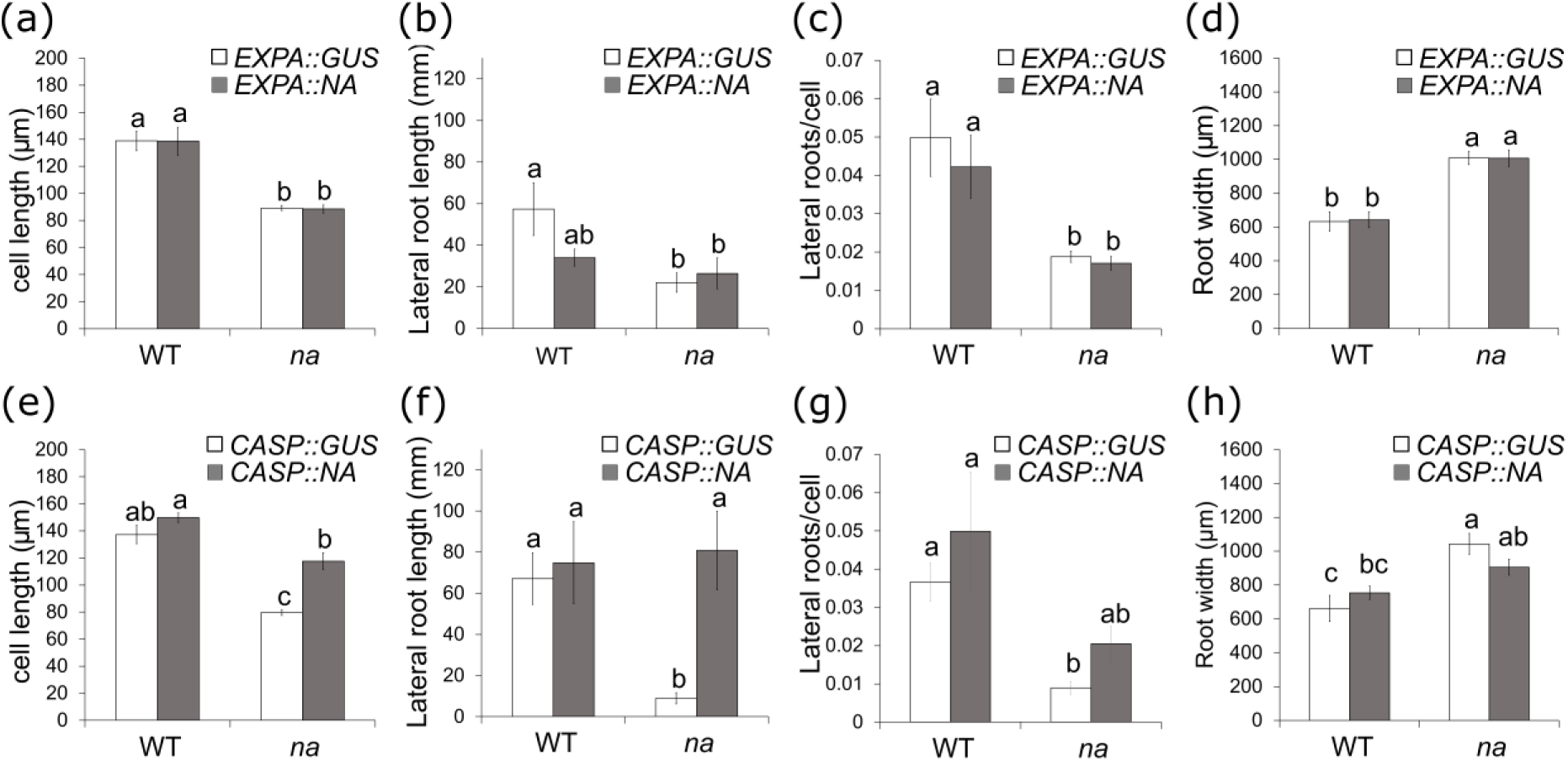
Root phenotype in endodermal (*CASP*) and epidermal (*EXPA*) *NA* complementation experiments in WT and the *na* mutant lines. (a-d) Quantitative evaluation of the root phenotype for the epidermal *NA* complementation experiment: cortical cell length (a), lateral root length (b), number of lateral roots (c) and root width (d). (e-f) Quantitative evaluation of the root phenotype of endodermal *NA* complementation experiment: cell length (e), lateral root length (f), number of lateral roots (g) and root width (h). Values are mean and bars represent the standard error. Grouping letters were obtained using one way ANOVA with Tukey *post hoc* test. Values with different letters are significantly different (P<0.05), (n = 5-12).

### Examining the correlations between gibberellin content and nodulation

The pea mutant *sln* offers an alternative way to evaluate the impact of the site of bioactive gibberellin production on infection and nodule development in pea. The *sln* mutant is disrupted in gibberellin catabolism gene *PsGA2ox-1* which encodes an enzyme that breaks down the bioactive GA_1_ and its immediate precursor GA_20_ (Martin *et al*., 1999). As *PsGA2ox1* is primarily expressed in the developing seed, the *sln* mutation leads to a build-up of bioactive GA_1_ in seedling shoot and root tissue. This effect is not observed in tissue that develops after the seedling stage due to the activity of an additional *Ps*GA2-oxidase enzyme in more mature vegetative pea tissue (Lester *et al*., 1999). Both the number of infections and number of nodules were significantly reduced in the top 40mm of the *sln* mutant root compared to wild type, although if nodules did develop they were of similar size to wild type as previously observed (Fig S3; Ferguson *et al*., 2005). This is consistent with previous results in pea *della* mutants, where increased gibberellin signalling suppressed infection and subsequent nodule number but did not impact on nodule size (Ferguson *et al*., 2011; McAdam *et al*., 2018) . However, in the tap root more distal from the cotyledons (40-140mm below the cotyledons) that developed post-seedling stage there was no significant change in infection or nodule number in the *sln* mutant plants compared with wild type (Fig. S3)

Previous studies have found that bioactive gibberellin levels are elevated in *Lotus* nodule tissue compared to uninfected roots (Akamatsu *et al*., 2021) and a gibberellin biosensor is active in the nodule meristem of soybean nodules (Chu *et al*., 2021). To determine the levels of gibberellins early in infection, gibberellins were monitored in the infection zone of wild type and supernodulating *Psnark* mutants. *Psnark* mutants have a defect in a peptide signalling protein important in autoregulation of nodulation (Sagan & Duc, 1996; Krusell *et al*., 2002). Plants were examined 2 dpi, just prior to the development of infection threads to enable us to examine if changes in gibberellin levels precede the elevated infection that develops in *Psnark* mutants comparted to wild type (Fig. S4a). No significant difference in the levels of the bioactive gibberellin (GA_1_) or key gibberellin precursors (GA_19_ and GA_20_) or gibberellin catabolite (GA_8_) were observed between wild type and *Psnark* mutant roots 2 dpi. This is consistent with previous findings that did not find any significant changes in the expression of gibberellin biosynthesis or catabolism genes in response to inoculation in wild type peas (McAdam *et al*., 2018) .

### Interaction of gibberellins and cytokinin during nodulation

Data presented above, and in previous studies, indicate gibberellins play distinct spatial roles in the root, suppressing infection in the epidermis and promoting nodule organogenesis and nodule development in the inner root layers. Interestingly, a somewhat similar role has been proposed for cytokinin (Gamas *et al*., 2017). Indeed, we found that while application of BAP to wild type pea had no effect on the number of normal infection threads, it did lead to the development of highly branched infection threads and suppressed the number of nodules (Fig. S5); the latter two phenotypes are also observed in gibberellin-deficient *na* mutant plants (e. g. Fig. 1, 2 and 4).

**Fig. 4.**
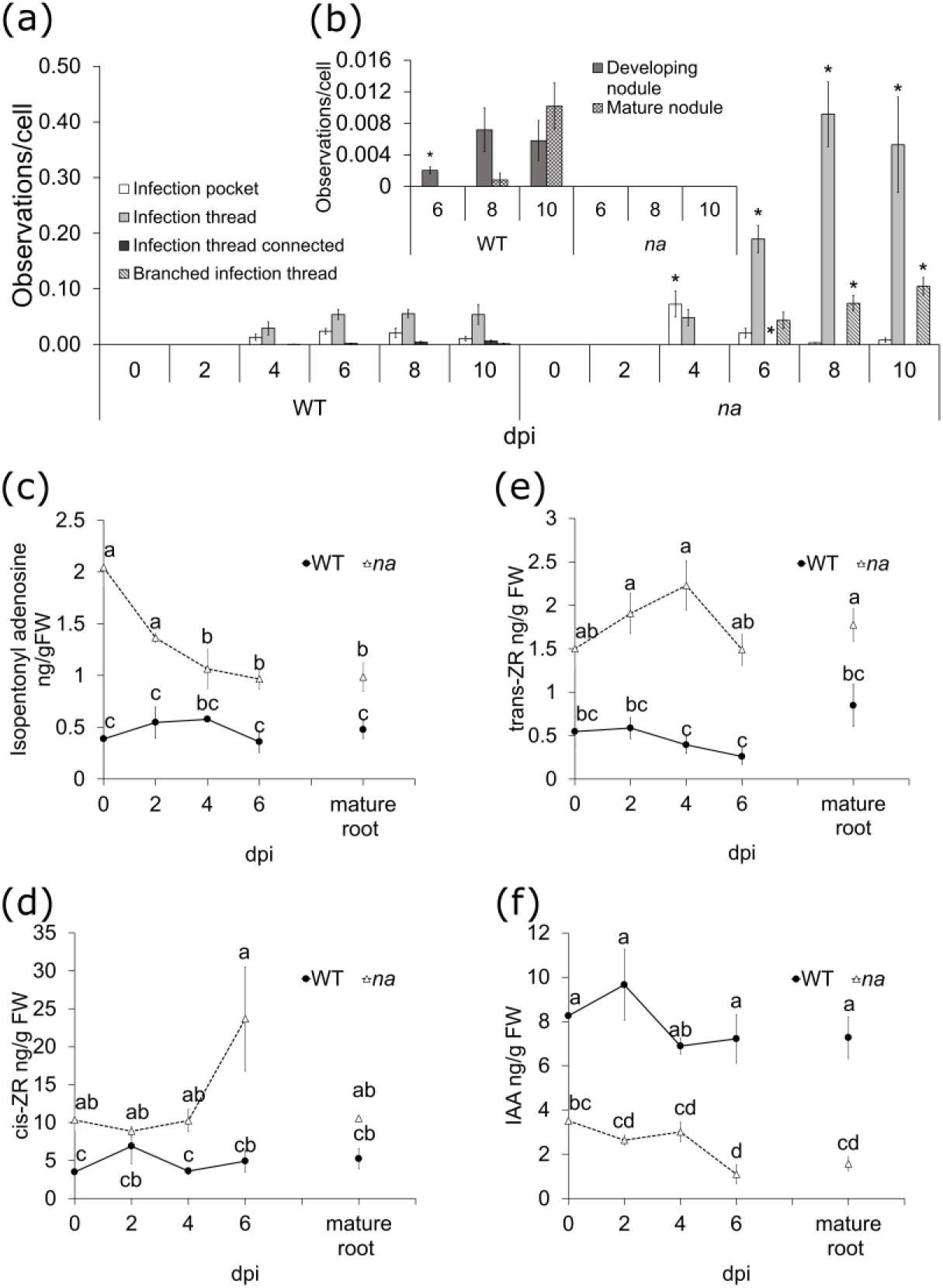
Nodulation and hormone dynamics in WT and the *na* mutant. (a-b) number of structures per cell 0, 2, 4, 6, 8 and 10 days post inoculation (dpi). Infection structures (a) and nodules (b) in WT and the *na* mutant line. (c-f) Cytokinin and indol-3-acetic acid (IAA) levels in the infection zone of WT and *na* mutants at 0,2,4 and 6 dpi and in mature roots 6 days post inoculation. Values are mean and bars represent the standard error. For (a,b) asterisks represent values significantly different between WT and the *na* mutant using t-test (* p<0.05), (n = 3). For (c-f) grouping letters were obtained using one way ANOVA with Tukey *post hoc* test or Kruskal for non-parametric data and values with different letters are significantly different (P<0.05), (n = 4).

To examine the potential interactions between gibberellins and cytokinin, several approaches were taken, including cytokinin quantification, the use of cytokinin biosensors and hormone application. Firstly, cytokinin levels during infection and nodule development were examined in a time course experiment with wild type and *na* non-transgenic plants. To establish when infection and nodule development occur in these genotypes, infection and nodule development was monitored every 2 dpi (Fig. 4a,b). The first infection events (the formation of infection pockets and infection threads in the epidermis) was observed in both wild type and *na* mutants 4 dpi. By 4 dpi, *na* mutant plants had a significant 5-fold increase in the number of infection pockets compared to wild type (Fig. 4a). From 6 to 10 dpi, *na* mutant plants had significantly more infection events, including branched infection threads in the cortex, compared to wildtype (Fig. 4a). Developing nodules were not observed until 6 dpi in wild type and mature nodules were present 8 dpi; in contrast no developing or mature nodules were observed in *na* mutants during this time course (Fig. 4b). Three cytokinin molecules could be reliably detected in pea root tissue; the precursor isopentonyl adenosine and the bioactive cytokinins cis- and trans-zeatin riboside (ZR) (Fig. 4c-e). Prior to inoculation, both isopentonyl adenosine and cis-ZR were significantly more abundant in the infection zone of *na* mutant plants compared to wild type (Fig. 4c,d). In wild type, we found no significant change in the levels of any cytokinin in response to inoculation in the infection zone, possibly because infection events occur in only a small number of cells or that cytokinin production is not activated by inoculation. In contrast, *na* mutant roots displayed a significant reduction in isopentonyl adenosine concentration 4 dpi, which was sustained at 6 dpi in the infection zone and in mature root tissue, compared to uninoculated *na* plants (Fig. 4c). However, isopentonyl adenosine concentration was still significantly higher 2 and 6 dpi in *na* mutants plants compared to wild type plants (Fig. 4c). An increase in cis-ZR was observed in the infection zone but not the mature root of *na* mutant plants 6 dpi compared to wild type 6 dpi (Fig. 4d). No significant changes in trans-ZR were observed in *na* mutant plants in response to inoculation, although trans-ZR levels remained significantly higher in all *na* mutant tissue from 2-6 dpi compared with wild type plants (Fig. 4e).

Cytokinin levels were also monitored in the transgenic experiments outlined above with epidermal and endodermal expression of the wild type *NA* gene in wild type and *na* mutants (Table S1). It is important to note that the transgenic tissues harvested for hormone analysis were whole roots that contained zones at different stages of infection, nodule development and root development 3 weeks post inoculation, which makes it hard to identify specific hormonal changes. The level of cytokinins in *na* mutants in the control, the epidermal and endodermal complementation roots was not markedly different to those from comparable WT roots (Table S1).

In order to visually examine cytokinin response during infection and nodule development in roots with altered gibberellin levels, the cytokinin responsive *TCSn::GUS* construct was expressed in gibberellin-deficient *na* mutants and wild type roots (Fig. 5). In order to capture the pattern of cytokinin response in *na* and wild type roots in relation to infection and nodule development, *TCSn::GUS* staining associated with infection threads not connected to a nodule, branched infection threads and infection threads connected to a nodule primordium were recorded. In wild type roots, the vast majority of infection threads in the epidermis not connected to a nodule displayed no *TCSn::GUS* staining (IT GUS-; Fig. 5c,e). However, *TCSn::GUS* staining was seen in almost all wild type infection threads connected to nodule primordium (IT connected GUS+; Fig. 5c, f). In addition, in wild type plants *TCSn::GUS* staining was observed in the whole developing nodule and the meristem of mature nodules (Fig. 5g,h). In stark contrast, in *na* mutants roots approximately 50% of normal and branched infection threads not connected to nodules displayed strong *TCSn::GUS* activity (IT GUS+; Fig. 5c, j). Given the large numbers of infection events in *na* mutants (approximately 10-fold increase in infection events compared to wild type: Fig. 5d) the young roots of *na* mutants had large patches of *TCSn::GUS* staining (Fig. 5b, j). This *TCSn::GUS* staining associated with branched infection threads of *na* extended through the cortex of *na* mutants but was not associated with any cell divisions or differentiation (Fig. 5k). In the very small number of developing nodules that did occur on *na* mutant roots in this experiment, there was associated *TCSn::GUS* staining; recorded as infection threads connected to nodule primordium (IT connected GUS+; 1% of all *na* infection threads; Fig. 5i).

**Fig. 5.**
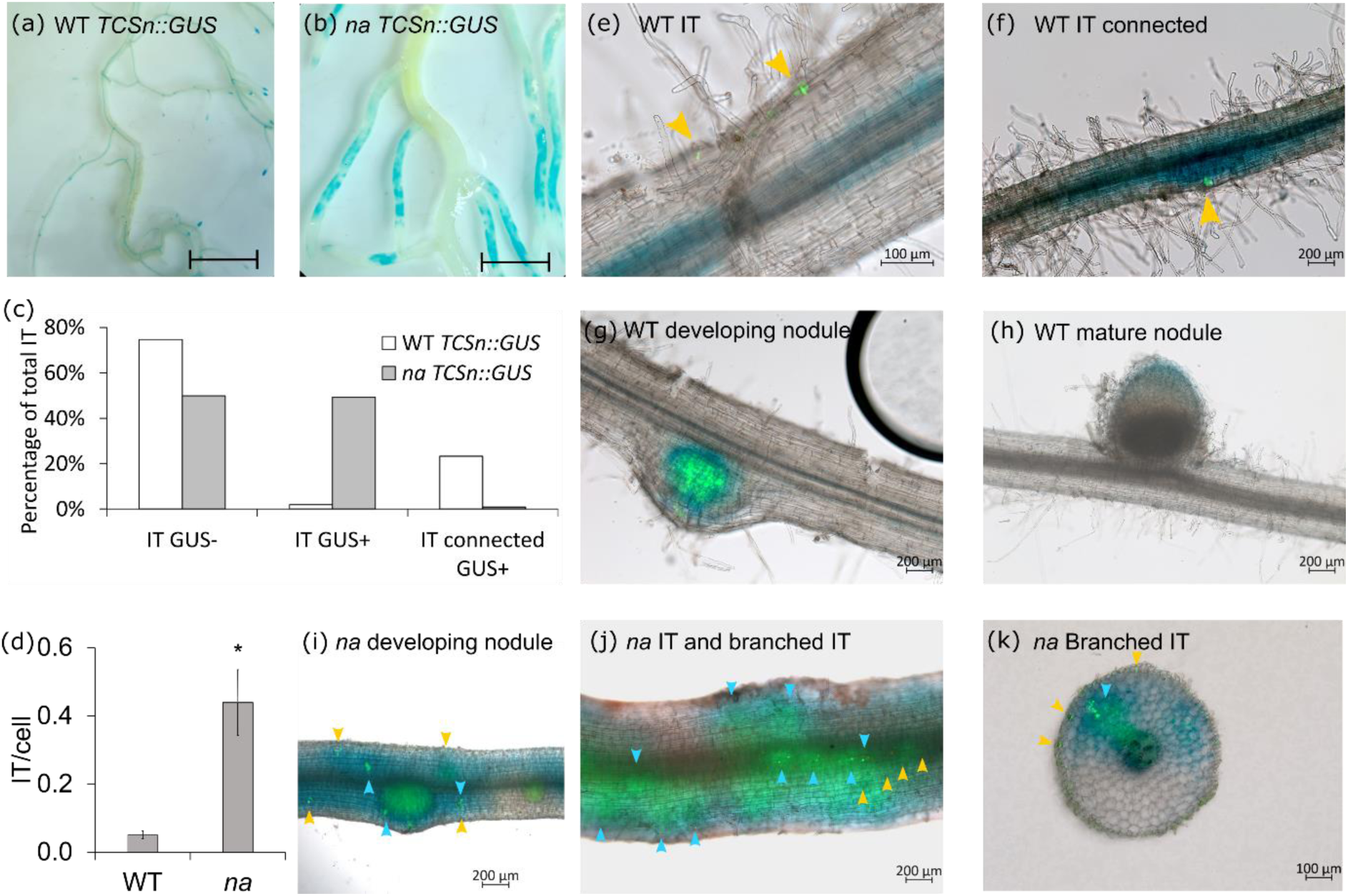
Cytokinin responsive promoter *TCSn* activity in WT and *na* mutants. Blue colour indicates GUS activity. (a, b) *TCSn::GUS* activity in roots 4 weeks after inoculation with rhizobia in (a) WT and (b) *na* mutant, scale bar is 1cm. (c) The percentage of total infection threads with GUS (IT GUS+), IT without GUS activity (IT GUS-) and IT connected with nodule primordia with GUS activity (IT connected GUS+). WT and *na* mutants showed a significant interaction between genotype and GUS activity (2x2 chi square test, P<0.001). (d) Number of IT per cell. Asterisks represent values significantly different between WT and the *na* mutant using t-test (* p<0.05), (n = 4-12) (e-g, i-k) Merged channels of fluorescent images showing rhizobia in green and bright field in WT and *na* mutant roots expressing *TCSn::GUS,* normal and branched infection threads are indicated with yellow and blue arrowheads respectively. Pictures of representative structures in WT roots: IT (e), IT connected to a nodule primordium (f), developing nodule (g) and mature nodule (h). Representative pictures in *na* mutant for developing nodules (i) and normal and branched IT in a longitudinal view (j) and a cross section (k) of the root.

The induction of nodule-like structures by cytokinin appears to be a legume-specific response and is a powerful tool to examine the role of this hormone in nodule organogenesis (Gauthier-Coles *et al*., 2019). However, in pea cytokinin application leads to excessive root swelling, making it very difficult to distinguish any possible cytokinin-induced nodule-like structures (data not shown; Gauthier-Coles *et al*., 2019). Therefore, soybean was used to test whether gibberellins are required for cytokinin to induce nodule-like structures (Fig. S6). The application of the synthetic cytokinin BAP induced the production of nodule-like structures in soybean. Gibberellin appears to be required for the development of these organs, as the number of pseudonodules was significantly increased in the presence of the gibberellin, GA_3_ (Fig. S6a) and was reduced when the gibberellin biosynthesis inhibitor PAC was applied (Fig. S6b).

### Interaction of gibberellins and auxin during nodulation

Auxin biosensors have been powerful tools to examine the role of auxin during nodulation. We examined the possible interaction between auxin and gibberellin by generating a *DR5::GUS*-tagged *na* mutant and corresponding wild type and examining the pattern of *DR5::GUS* staining during nodulation (Fig. 6). We did not detect *DR5::GUS* staining associated with infection threads in WT or *na* roots, including with branched infection threads seen in the cortex of *na* mutant roots (Fig. 6 a, b, g, h). In wild type roots *DR5::GUS* staining was observed in root tips, during early nodule development in the nodule primordium and became restricted to the nodule meristem and vascular bundles in mature nodules (Fig. 6 c-f, m). In gibberellin-deficient *na* mutants *DR5::GUS* staining was also observed in the root tips (Fig. 6. n). Importantly, we found an absence of *DR5::GUS* staining in nodule primordium of *na* mutants (Fig. 6 i, j). The very few nodules that did develop past this stage in *na* mutant roots displayed *DR5::GUS* staining associated with the nodule meristem and vascular tissue as seen in wild type (Fig. 6 k-l).

**Fig 6.**
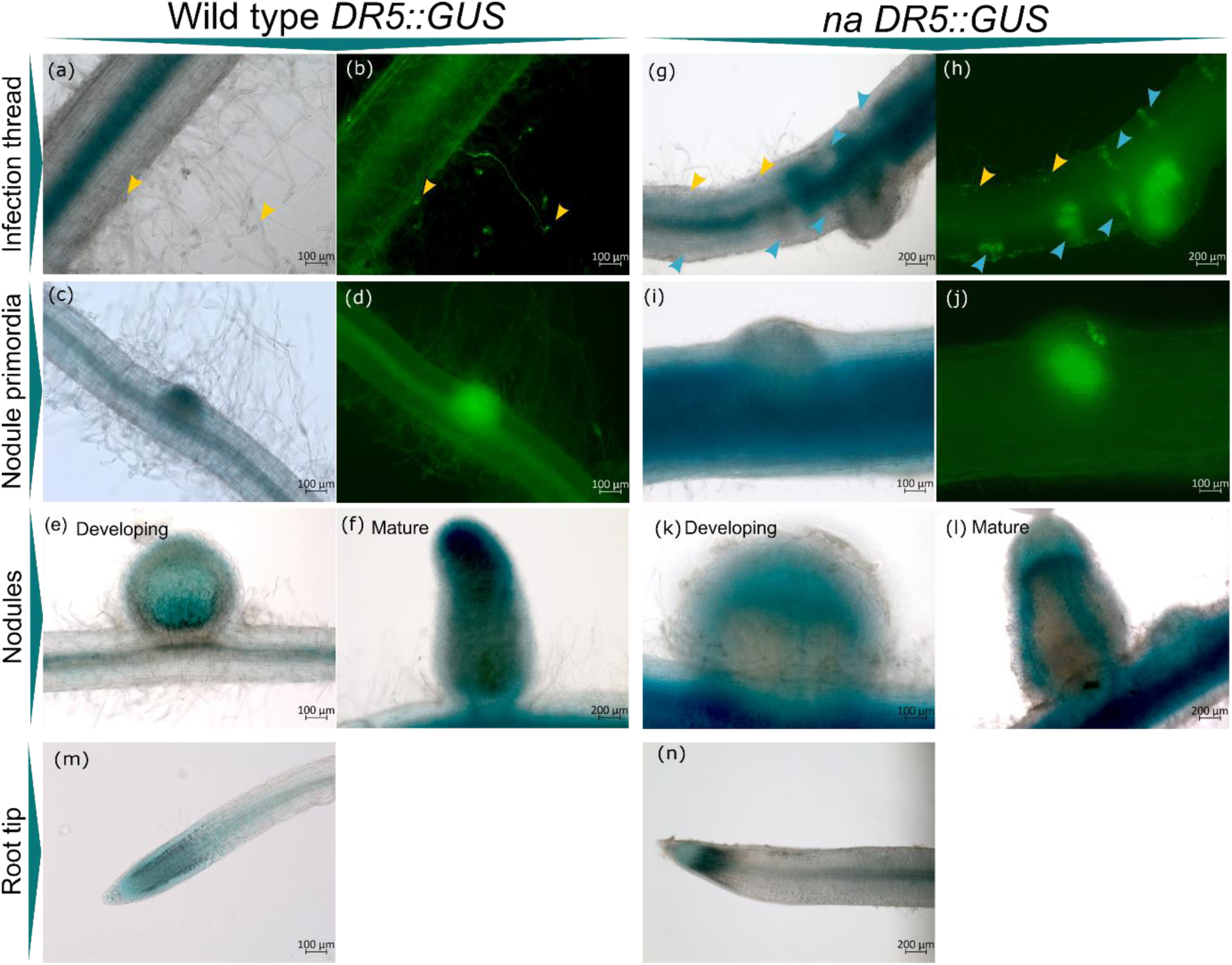
Auxin responsive promoter *DR5::GUS* activity in WT and *na* mutant lines four weeks post inoculation with rhizobia. Blue colour indicates GUS activity and green fluorescence indicates rhizobia. (a-f) Pictures of representative structures in WT roots in brightfield (a, c, e and f) and corresponding blue channel fluorescence (b and d), (a-b) infection threads (yellow arrowheads), (c-d) nodule primordia, (e) developing nodule and (f) mature nodule. (g-l) Pictures of representative structures in *na* mutant roots in bright field (g, i, k and l) and corresponding blue channel fluorescence (h and j), (g-h) normal infection threads (yellow arrowheads) and branched infections (blue arrowheads), (i-j) nodule primordia, (k) developing nodule and (l) mature nodule. (m-n) *DR5* response in WT (m) and *na* (n) root tips.

Auxin levels were also monitored in the time course experiment with wild type and *na* mutants outlined above (Fig. 4f). Overall, *na* mutant roots had significantly less auxin content than wild type roots, and this is consistent with reduced auxin content previously seen in *na* mutant root tips (Silva & Davies, 2007). There was no significant change in auxin levels in response to inoculation in *na* mutant plants. In wild type the only small but significant changes seen in IAA was a small decrease in IAA levels in the infection zone 6 dpi compared to pre-inoculation.

Auxin transport is reported to play a key role in nodule organ formation (Ng *et al*., 2015), and in many species application of auxin transport inhibitors can induce nodule-like structures in the absence of rhizobia (Rightmyer & Long, 2011). To further examine the interactions between auxin and gibberellins, the auxin transport inhibitor NPA was applied to sterile wild type and *na* mutant roots (Fig. S7). NPA induced a similar number of nodule-like structures in both wild type and *na* mutants, indicating that gibberellins are not required or act upstream of auxin in the development of these structures.

## Discussion

In this study, we have identified a fundamental role for GA, produced in the endodermis, in the regulation of nodulation. In the absence of GA, pea plants fail to develop mature functional nodules, and the nodule primordia that do form exhibit impaired growth and reduced ability to host rhizobia (Fig. 1-2). However, the restoration of GA biosynthesis in the endodermis of *na* mutants rescues its ability to form mature functional nodules (Fig. 2 i, m). These nodules exhibit a characteristic pink colour, are filled with bacteria, and reach a size comparable to wild-type nodules (Fig. 2f, g-i). Interestingly, while GA does not appear to be essential for nodule initiation (Fig. 2l), the progression of nodule development does require GA, indicating that GA likely acts as a checkpoint mediating the transition from nodule primordia formation to full maturation. Furthermore, our investigation reveals that the presence of GA in the epidermis alone suppresses infection but fails to restore nodule development (Fig. 1f, m), suggesting that GA has limited mobility from the epidermis to the endodermis during nodulation and implying tissue-specific actions of GA in the root. It is also clear that the nodule number and size is not an indirect effect of shoot size, as dwarf shoots were present on *na CASP::NA* plants that developed many mature sized nodules (Fig. S2e). Our findings indicate that GA is required specifically in the inner root layers to promote the progression of nodule formation (Fig. 7) and this is consistent with research across other legumes demonstrating a positive role of GA in nodule organogenesis (Ferguson *et al*., 2005; Kim *et al*., 2019; Chu *et al*., 2021; Gao *et al*., 2023).

**Fig. 7.**
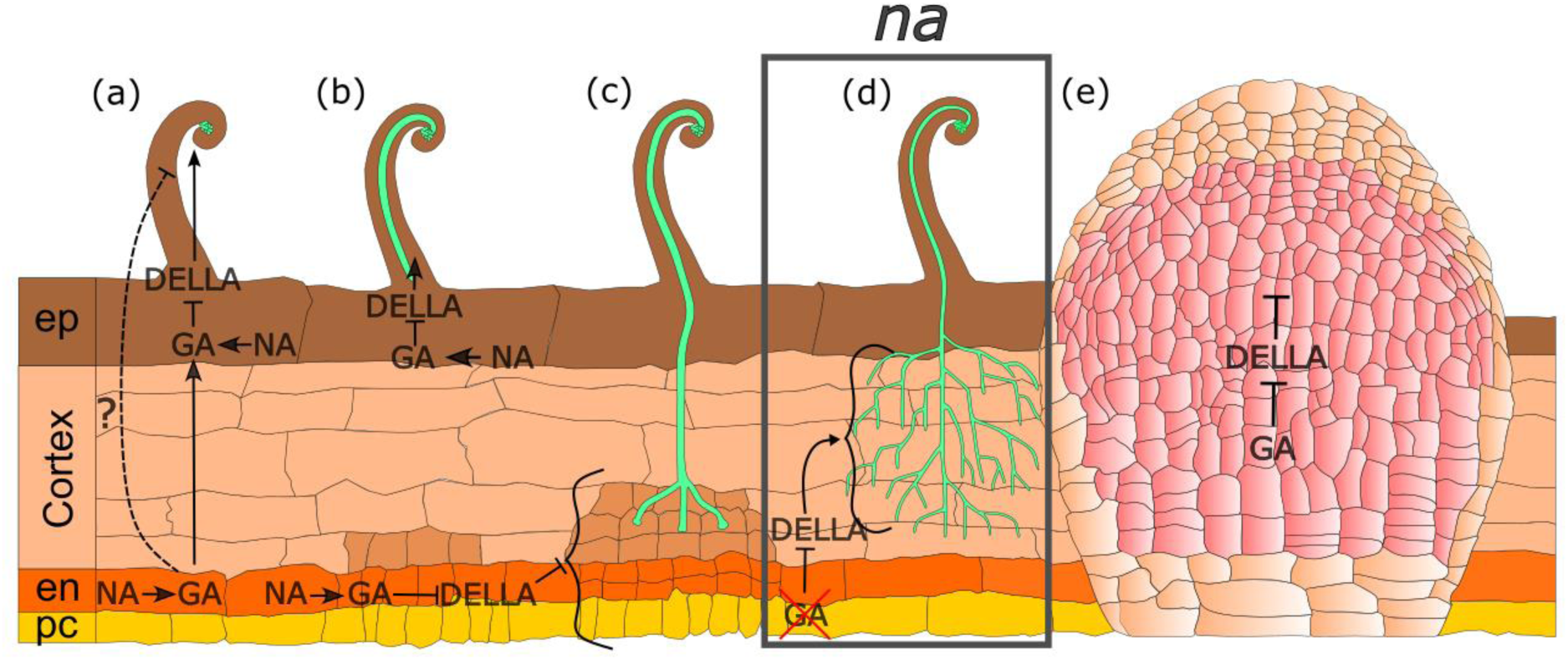
Model of the role of gibberellin (GA) via DELLA degradation in regulating rhizobial infection and nodule organogenesis in pea. (a) GA inhibits the earliest stages of infection at the epidermis including the formation of infection pockets. Infection thread (IT) (green) formation is inhibited by GA produced in the epidermis or endodermis. GA may act directly or via another mobile element from the endodermis to limit IT formation. (b) GA limits infection thread branching in the cortex. (c) GA in the endodermis promotes nodule development and maturation. (d) In the absence of GA *(na* mutants, box), infection threads branch in the cortex and fail to elicit cell division and colonisation of the nodule primordia. (e) GA promotes nodule maturation and function. ep: epidermis (brown), en: endodermis (orange), pc: pericycle (yellow), nodule cells infected with rhizobia are pink.

Our results conclusively demonstrate that GA restricts the formation of infection threads in the epidermis (Fig. 7a-b). A reduction in GA levels, as seen in *na* mutants, increase the very earliest stages of infection in the epidermis, infection pocket formation (Fig. 4a) and elevated levels of GA, as seen in the top section of the tap root of *sln* mutant, decreases infection (Fig. S3). Intriguingly, GA produced in the epidermis, or the endodermis could suppress infection, indicating GA or another mobile element that can suppress infection moves from the inner root layers to the epidermis. This signal does not appear to be cytokinin, as cytokinin levels and response is actually over-activated in *na* roots which display very high infection events (Fig. 4, 5). Given that nodules may produce gibberellins (Akamatsu *et al*., 2021), this effect may be a feedback mechanism to control infection after nodulation.

GA also plays a role in regulating the progression and branching of infection threads through the cortex (Fig. 2k, 7d). In *na* mutants, approximately 17% of the total infection threads become highly branched in the cortex and fail to trigger cell division and colonisation of a nodule primordia (Fig. 1b, k, 2b, k). This abnormal branching of infection threads is absent in wild type plants, strongly suggesting that it is a consequence of GA deficiency. Upon the restoration of GA in the endodermis, a significant reduction in branched infection threads was observed, indicating that GA in the inner root layers partially restricts infection thread branching (Fig. 7d). This significant reduction was not observed in epidermal complementation (Fig. 1k). Studies suggest that there are signals directing infection threads growth inward to colonise the nodule primordia, where branching is normally triggered (Monahan-Giovanelli *et al*., 2006). However, the precise nature of this signal remains unclear. It is possible that changes in polarity and growth direction are regulated by a mobile signal originating from the inner layers of the root. One potential mechanism could involve the presence of a GA gradient, acting as a signal to guide the progression of infection threads. This guidance might be mediated through the degradation of DELLA proteins or the restriction of cytokinin activity during infection thread growth discussed below.

We found that GA acts both up and downstream of CK during nodulation (Fig. 8). GA appears to negatively regulate CK level and response, as we observed elevated levels of CK (Fig. 4c-e) and a pronounced overactivation of the CK biosensor surrounding branched infection threads in GA-deficient *na* roots (Fig. 5).This suggests GA restricts CK response in the cortex and the fact that application of exogenous CK in pea also leads to infection thread branching (Fig. S5), indicates that GA may suppress infection thread progression and branching by suppressing CK in the cortex. CK signalling is reported to play a pivotal role in activating the nodulation pathway and initiating cell division in the cortex (Heckmann *et al*., 2011). This CK activation during nodulation is dependent on DELLA signalling and inhibited by the presence of GA (Fonouni-Farde *et al*., 2017). Our findings suggest that GA acts upstream of cytokinin biosynthesis and/or response to enforce this checkpoint regulating infection thread entry into the cortex. It also appears that GA may act downstream of CK activation to facilitate the progression of nodule development. This is supported by the fact that high doses of CK application result in reduced nodule formation in pea (Fig. S5), and that in soybean the induction of nodule-like structures by cytokinin depends on the presence of gibberellin (Fig. S6).

**Fig. 8.**
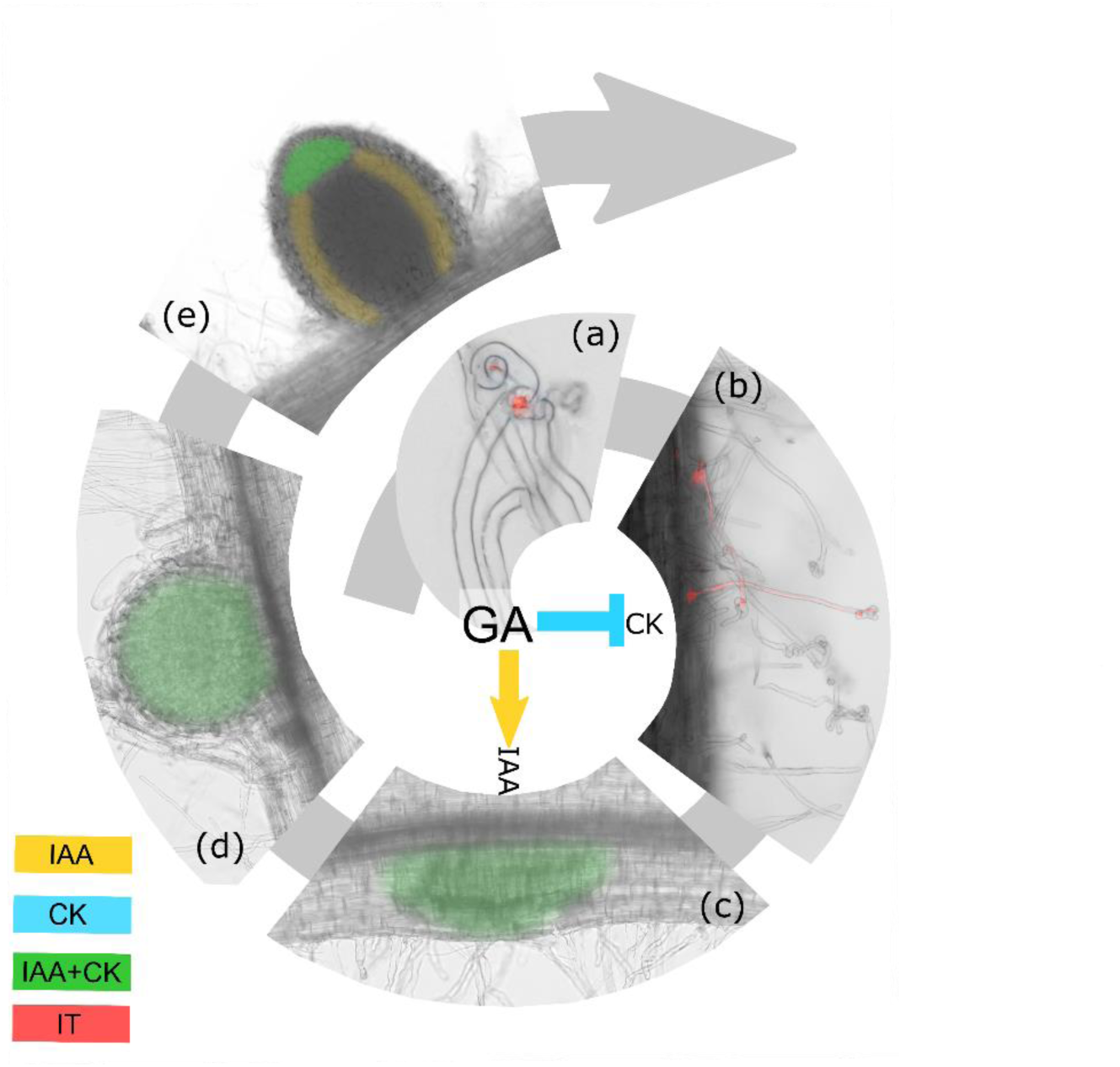
Auxin (IAA) and cytokinin (CK) response patterns during nodulation and interaction with gibberellin (GA) in pea. (a-b) During infection pocket and infection thread formation IAA and CK-responsive promoter activity is not detected in wild type. Low GA levels (*na* mutants) leads to branched infection threads developing in the epidermis and cortex that are associated with ectopic activation of a CK-responsive promoter. (c-d) CK and IAA response is activated in the dividing cells of the nodule primordia before bacteria reach the inner cortex (c). This suggests that there is a signal from the epidermis that activates cell division and IAA/CK response. GA is required for activation of the IAA response in the nodule primordia. (d). CK and IAA response occurs throughout developing nodules. (e) In mature nodules CK response is restricted to the nodule meristem and IAA response occurs in the nodule vasculature and meristem.

Our results indicate GA may activate auxin response to promote nodule primordia formation but not necessarily later nodule development (Fig. 8). In WT and *na* plants, *DR5* promoter activity was not observed in association with normal or branched infection threads (Fig. 6). However, in wild type plants *DR5* was activated early in nodule primordia formation, and this was not observed in *na* nodule primordia. This lack of auxin response could be attributed directly to the absence of GA or may be due to the high levels of CK in these *na* roots. The majority of *na* nodules do not mature fully, although the few nodules that were observed to develop on *na* mutants displayed a similar pattern of auxin response in the meristem and vasculature as seen in wild type. Intriguingly, *na* mutants could form auxin-induced pseudonodules upon treatment with the auxin transport inhibitor NPA, supporting the notion that GA is not required downstream of auxin during nodule development (Fig. S7). It is interesting to note that for both auxin and CK, hormone measurements of whole root tissues were not able to provide insights into precise spatially regulated hormone interactions.

GA is required in the endodermis for root development. The absence of GA has a significant impact on cell length, cell width, and development of lateral roots (Fig. 3). In our study, only endodermal complementation restored the root phenotype of GA-deficient lines (Fig. 3). This suggest that GA is required in inner root tissues for normal root development. This finding aligns with previous research conducted on *Arabidopsis*, which has demonstrated that GA metabolic enzymes are predominantly expressed in the endodermis, facilitating cell elongation and maintaining appropriate meristem size (Ubeda-Tomás *et al*., 2009; Barker *et al*., 2021). The failure in root phenotype recovery in epidermal complementation further support the idea of a lack of GA movement from the epidermis to the inner root tissues. Whether GA exhibits limited mobility in pea roots or if the amount of GA produced is insufficient for transport remains to be determined. The simultaneous restoration of nodulation and root phenotypes by endodermal GA (Fig. 2, 3) suggests the presence of a shared mechanism governing both developmental processes. Notably, numerous proteins crucial for root organogenesis are activated during nodulation, implying a potential evolutionary link between nodule and lateral root development programs (Soyano *et al*., 2021). GA may play a mediating role in activating organogenesis programs shared between root and nodule development, acting as a crucial step in coordinating both processes.

Our results demonstrate that GA plays dual and opposing roles in specific cell layers during infection and nodule organogenesis. We demonstrate that GA limits infection in the epidermis either by direct action or the potential involvement of another mobile signal(s) that can move from the endodermis to the epidermis. Moreover, GA is necessary for guiding infection threads towards nodule primordia by limiting excessive IT branching. We demonstrate the requirement of GA in the endodermis for promoting nodule and root development and that GA coordinates the response of cytokinin and auxin, leading to successful nodulation. Novel methodologies, such as imaging mass spectrometry (Hu *et al*., 2021), are now needed to enable precision mapping of the spatial distribution of hormone levels within tissues during these complex developmental processes. Future studies may examine if the interaction between GA and nodulation specific elements such as NIN (Akamatsu *et al*., 2021; Gao *et al*., 2023) also follow precise spatial and temporal patterns and how this intersects with the root development program.

## Data availability

Accession numbers in NCBI database for *PsNANA* AF537321.1, *AtCASP* ptomoter HQ699533.1., *PsEXPA* promoter BankIt2740115 PsEXPA OR509285.

## Acknowledgements

We thank Ellen Bedell for technical assistance, Maria Soto (University of Granada) for the gift of GFP-labelled *Rhizobium*, Sandra Bensmihen (INRA, France) for the *pCAMBIA CASP* construct, Bruno Mueller (Zurich-Basel Plant Science Center, Department of Plant and Microbial Biology, UZH) for the *TCSn* promoter, and David Nichols (CSL, UTAS) for expert assistance with hormone analysis. We thank Michell Lang, Tracy Winterbottom, and Sarah Kane for assistance with plant husbandry.

## Author contributions

E.F. conceived the project. K.V., A.C., E.F., and P.M. performed the experiments and analysed the data. JBR provided substantial intellectual contribution. K.V, JBR and E.F wrote the manuscript with input from all authors.

