## Supplementary information for "Cell layer specific roles for hormones in root development: Gibberellins suppress infection thread progression, promote nodule and lateral root development in the endodermis and interact with auxin and cytokinin"

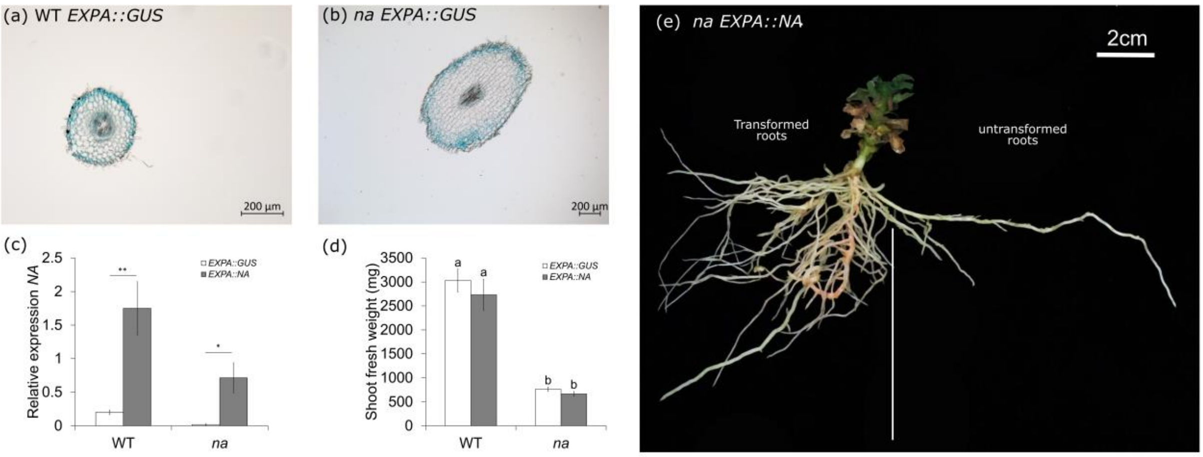


**Fig. S1** Characterization of the *PsEXPA* promoter in wild type (WT) and *na* mutant. (a,b) Tissue specific expression of *EXPA::GUS* construct (blue staining) in roots of WT and *na* mutants. (c) Relative expression of *NA* gene compared to *TFII* gene in WT and *na,* asterisks indicate statistical differences between pairs by t-test (* P<0.05, ** P<0.01) (n=5). (d) Shoot fresh weight measured at harvest for WT and the *na* mutant. Grouping letters were obtained using one way ANOVA with Tukey *post hoc* test and values with different letters are significantly different (P<0.05) (n=15-22). (e) Comparison of *EXPA::NA* transformed (left side) and untransformed (right side) roots in an *na* mutant plant. (c, d) Values are mean and bars represent the standard error.

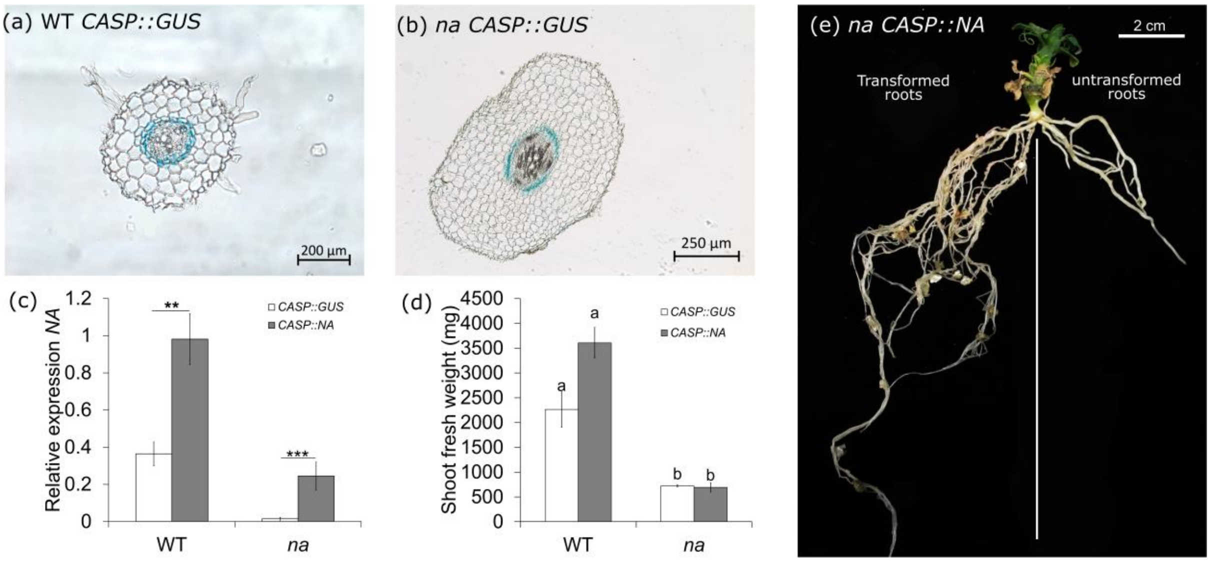


**Fig. S2** Characterization of *CASP* promoter in wild type (WT) and *na* mutant. (a,b) Tissue specific expression of *CASP::GUS* construct (blue staining) in roots of WT and *na* mutants. (c) Relative expression of *NA* gene compared to *TFII* gene in WT and *na,* asterisks indicate statistical differences between pairs by t-test (** P<0.01, *** P<0.001) (n=5). (d) Shoot fresh weight measured at harvest for WT and the *na* mutant. Grouping letters were obtained using one way ANOVA with Tukey *post hoc* test. Values with different letters are significantly different (P<0.05) (n=6-8). (e) Comparison of *CASP::NA* transformed (left side) and untransformed (right side) roots in *na* mutants. (c, d) Values are mean and bars represent the standard error.

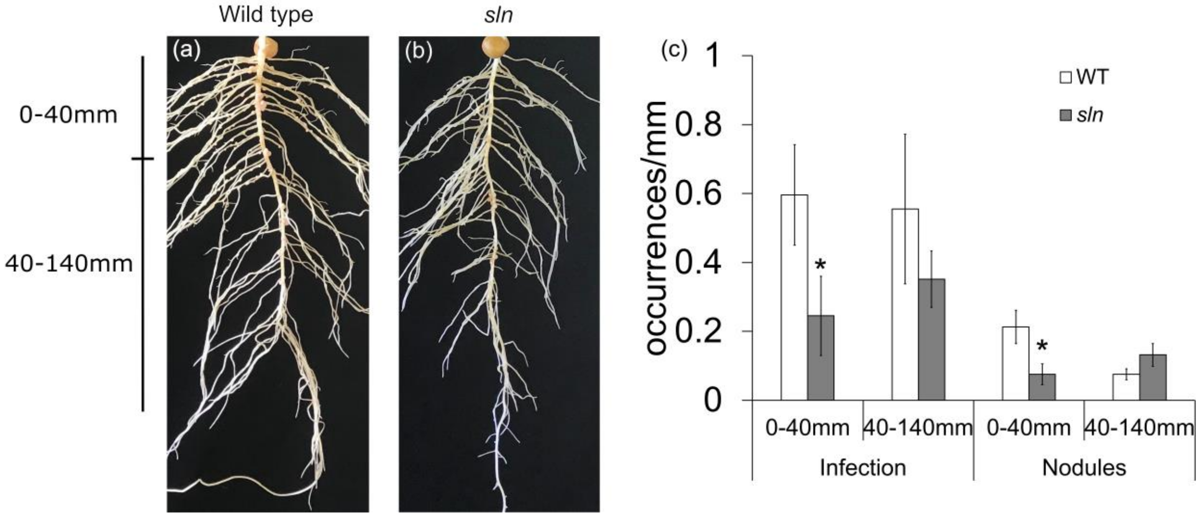


**Fig. S3.** Infection and nodule development in tap roots of wild type (WT) and *slender (sln)* mutant. (a-b) Photos of representative plants showing nodulation phenotype. (c) Number of infections and nodules per mm in the top 40mm of the taproot and from 40 to 140 mm of the tap root. Values are mean and bars represent the standard error. Asterisks indicates statistical differences between pairs by t-test (* P<0.05), (n = 6).

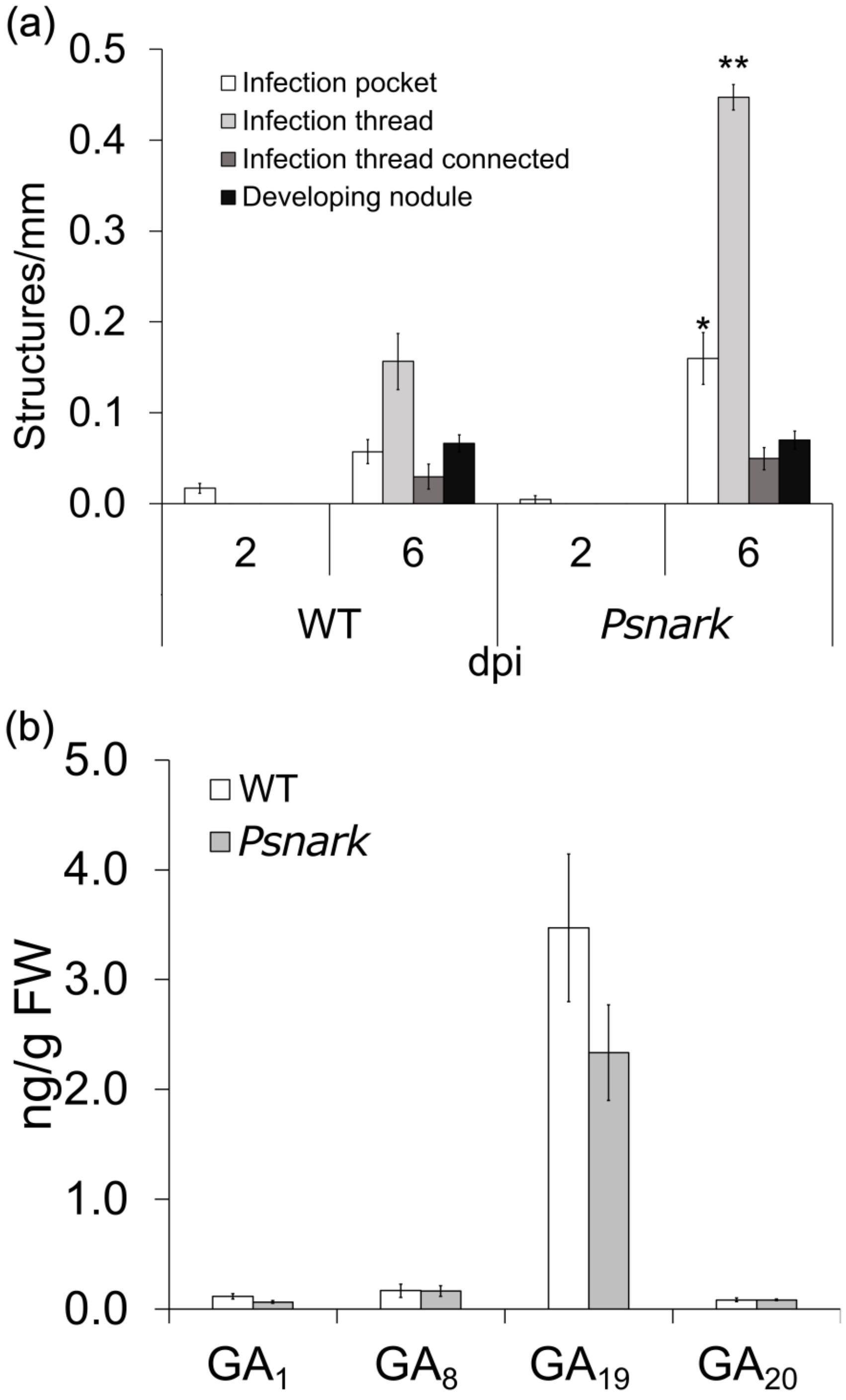


**Fig. S4.** Nodulation and gibberellins levels in wild type (WT) and *Psnark* mutant roots. (a) Number of infections and developing nodules per mm in plants 2 and 6 days post inoculation (dpi) (n = 6). (b) Gibberellin levels in the infection zone of roots 2 dpi (n = 3). Values are mean and bars represent the standard error. Asterisks indicates statistical differences between genotype pairs by t-test (*P<0.05, **P<0.01).

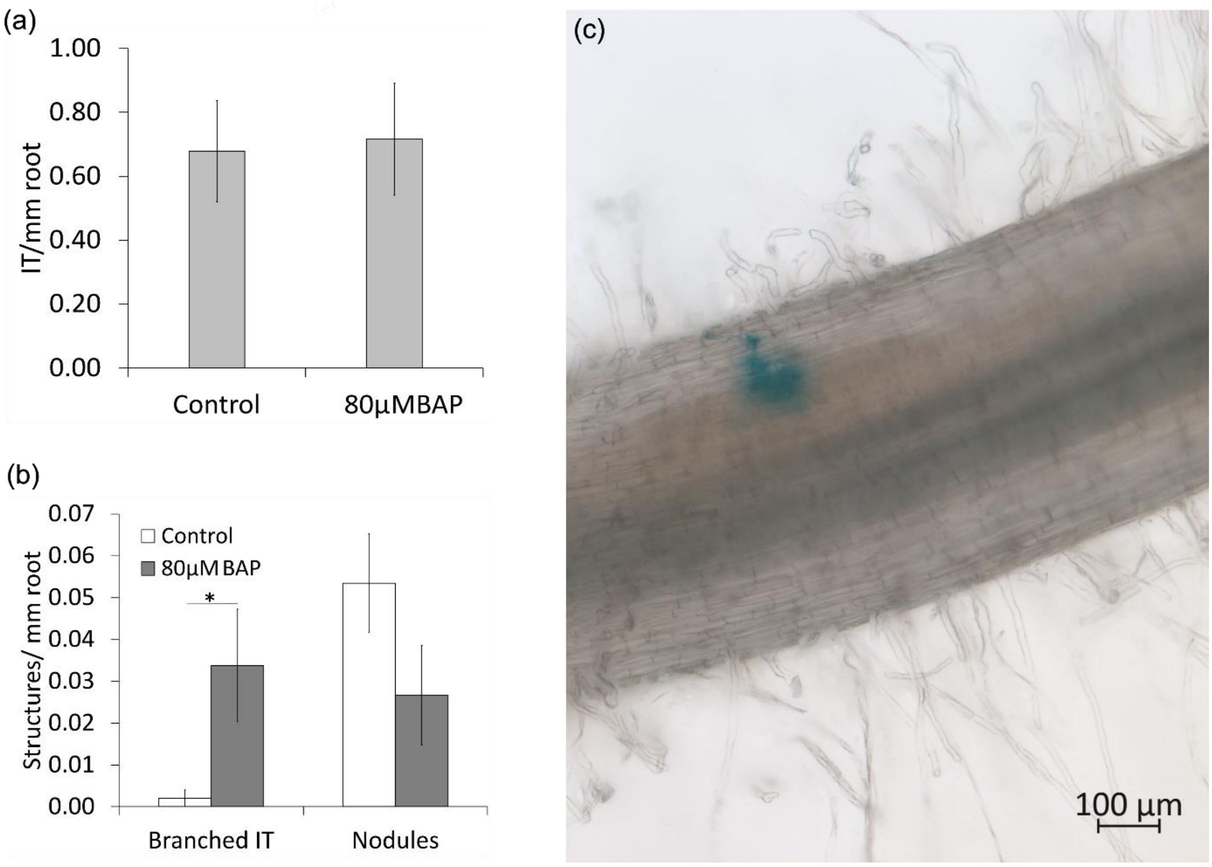


**Fig. S5.** Effect of 6-Benzulaminopurine (BAP) on nodulation in wild type (WT). Number of (a) infection threads (IT) per mm of root and (b) branched IT and nodules in control and plants treated with 80µM of BAP 3 weeks post inoculation. (c) Representative picture of a branched IT, rhizobia in blue. (a,b) Values are mean and bars represent the standard error and asterisks indicates statistical differences between pairs by t-test (*P<0.05), (n = 9-10).

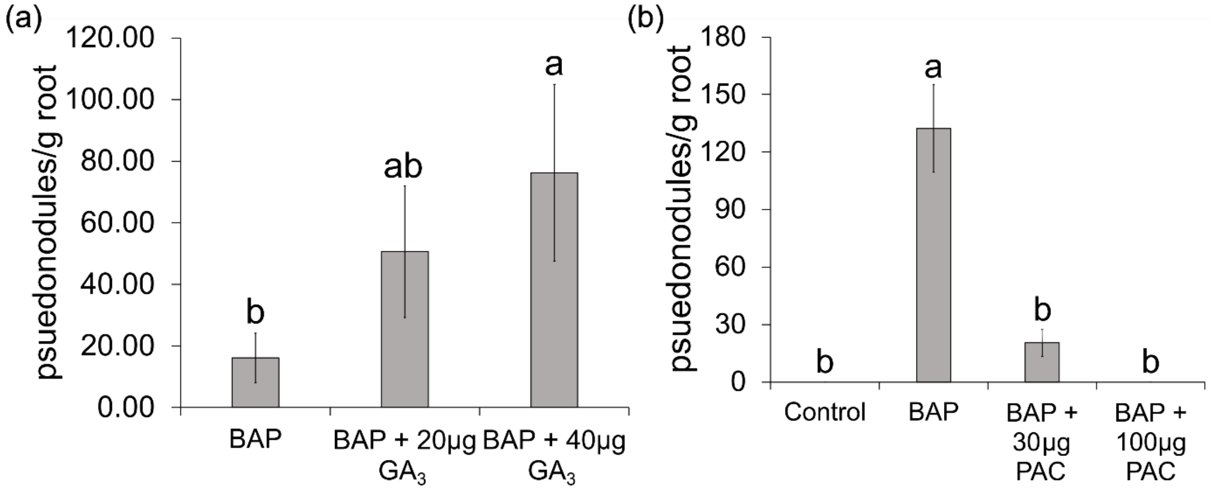

**Fig. S6**. Gibberellin is required for 6-Benzylaminopurine (BAP) induced pseudonodules in soybean. (a) Number of pseudonodules per gram of root dry weight in soybean plants treated with 20μM BAP and increasing amounts of $GA_3$. Grouping letters were obtained using one way ANOVA with Tukey *post hoc* test. Values with different letters are significantly different ($P<0.05$) ($n = 4$-12). (b) Number of pseudonodules per gram of root dry weight in soybean plants treated with 20 μM BAP and increasing amounts of GA biosynthesis inhibitor Paclobutrazol (PAC). Grouping letters were obtained using Kruskal-Wallis test and Bonferroni comparison. Values with different letters are significantly different ($P<0.05$) ($n= 9$-13). Values are mean and bars represent the standard error.

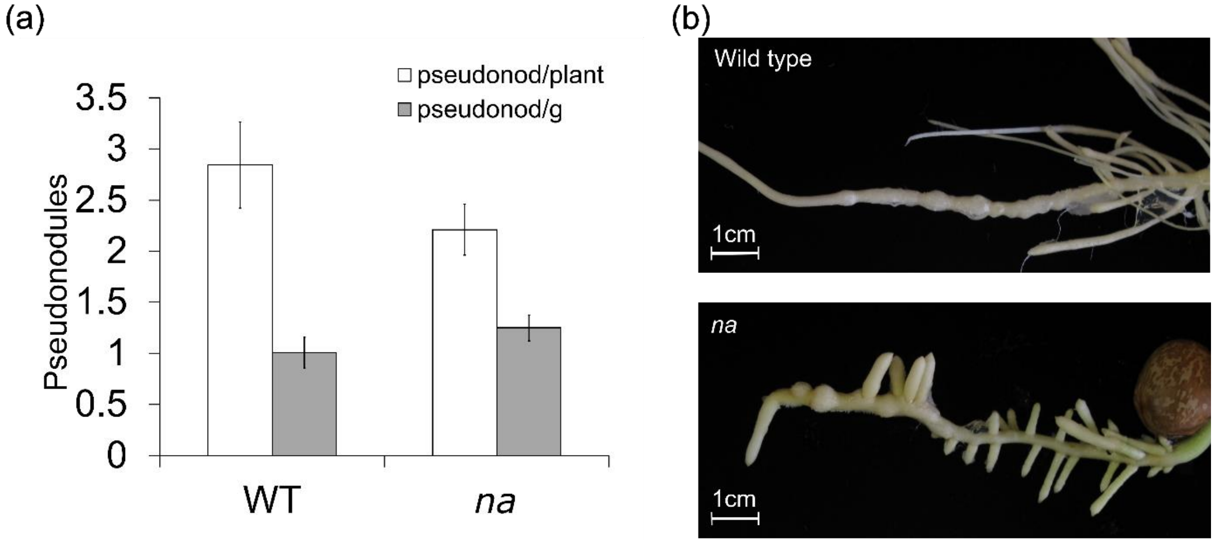


**Fig. S7.** Formation of pseudonodules in wild type (WT) and *na* plants after addition of auxin transport inhibitor naphthylphthalamic acid (NPA). (a) Number of nodule-like structures per plant and gram of root tissue, values are means, and bars represent the standard error. (b) Photos of a representative WT and *na* roots treated with NPA.

| **Genotype** | **Construct** | **trans-zeatin riboside** | **Isopentenyl adenine** | **Isopentenyl adenosine** | **cis-zeatin riboside** | **Indole acetic acid** |
|---|---|---|---|---|---|---|
| WT torsdag | *CASP::GUS* | 0.9114 | 0.1066 | 0.1738 | 0.6029 | 3.2051 |
| WT torsdag | *CASP::GUS* | 0.9690 | 0.1346 | 0.4127 | 0.8960 | 2.7355 |
| WT torsdag | *CASP::GUS* | 0.5830 | 0.0719 | 0.2165 | 0.3902 | 0.9295 |
| WT torsdag | *CASP::GUS* | 0.3755 | 0.0805 | 0.1927 | nd | 9.8540 |
| WT torsdag | *CASP::NA* | 0.2462 | 0.0724 | 0.2255 | 0.8472 | 44.2666 |
| WT torsdag | *CASP::NA* | 1.0282 | 0.0633 | 0.1366 | nd | 6.5950 |
| WT torsdag | *CASP::NA* | 4.8633 | 0.1984 | 0.5154 | 1.5882 | 9.8827 |

| | | | | | | |
|---|---|---|---|---|---|---|
| WT torsdag | *CASP::NA* | 4.7992 | 0.2236 | 0.5159 | 1.1567 | 26.8227 |
| *na* | *CASP::GUS* | 0.3577 | 0.0649 | 0.1915 | 1.7284 | 16.0494 |
| *na* | *CASP::GUS* | 0.6496 | 0.1218 | 0.3240 | nd | 12.0239 |
| *na* | *CASP::GUS* | 0.3475 | 0.1167 | 0.3523 | 2.6822 | 5.4520 |
| *na* | *CASP::GUS* | 0.2224 | 0.0811 | 0.1248 | 1.8772 | 10.7455 |
| *na* | *CASP::GUS* | 0.1317 | 0.0400 | 0.1130 | 2.4052 | 10.2214 |
| *na* | *CASP::NA* | 0.1094 | 0.0375 | 0.0970 | nd | 7.6398 |
| *na* | *CASP::NA* | 0.2671 | nd | nd | nd | 1.5986 |
| *na* | *CASP::NA* | 0.2399 | 0.0357 | 0.1382 | 0.4144 | 3.4053 |
| *na* | *CASP::NA* | 0.0552 | 0.0359 | nd | nd | 6.3951 |
| WT torsdag | *EXPA::GUS* | 1.0765 | 0.0294 | 0.4765 | 3.3088 | 58.5082 |
| WT torsdag | *EXPA::GUS* | 0.4872 | nd | 0.3533 | nd | 86.3785 |
| WT torsdag | *EXPA::GUS* | 1.1550 | nd | 0.2914 | nd | nd |
| WT torsdag | *EXPA::GUS* | nd | nd | 0.5073 | 2.5667 | 73.6098 |
| WT torsdag | *EXPA::GUS* | 1.1585 | nd | 0.3115 | 2.4281 | 99.9986 |
| WT torsdag | *EXPA::NA* | nd | 0.0187 | 0.2687 | nd | 22.5485 |
| WT torsdag | *EXPA::NA* | 4.8619 | 0.0865 | 1.1225 | nd | 54.3172 |
| WT torsdag | *EXPA::NA* | 2.0069 | nd | 0.8298 | nd | 88.0486 |
| WT torsdag | *EXPA::NA* | 1.1239 | nd | 0.4014 | nd | 133.3053 |
| WT torsdag | *EXPA::NA* | 0.3719 | 0.0302 | 0.1016 | nd | 31.2001 |
| *na* | *EXPA::GUS* | nd | 0.0434 | 0.3778 | 5.8540 | nd |
| *na* | *EXPA::GUS* | nd | 0.0235 | 0.3916 | 2.6424 | 32.0418 |
| *na* | *EXPA::GUS* | nd | nd | 0.3253 | 4.6940 | 34.9235 |
| *na* | *EXPA::GUS* | 0.1251 | nd | 0.0439 | 0.3372 | 2.3984 |
| *na* | *EXPA::GUS* | nd | 0.0018 | 0.0531 | 0.5897 | 9.8285 |
| *na* | *EXPA::NA* | nd | nd | 0.3247 | 4.8985 | 140.1994 |
| *na* | *EXPA::NA* | nd | nd | 0.0515 | 3.2655 | 26.1379 |
| *na* | *EXPA::NA* | 0.3685 | nd | 0.1670 | 2.1674 | 12.6997 |
| *na* | *EXPA::NA* | 0.4537 | 0.0776 | 0.3241 | nd | 56.9020 |
| *na* | *EXPA::NA* | 0.7033 | nd | 0.2616 | nd | 57.3273 |

**Table S1.** Cytokinin levels in whole transformed roots for the transformation experiments with wild type (WT) and *na* plants shown in Fig 1 and Fig 2. Quantification of cytokinins trans-zeatin riboside, isopentenyl adenine, isopentenyl adenosine and cis-zeatin riboside and indole-3-acetic acid in individual replicates. Other cytokinin species were not detected (nd).

| Target | Primers sequence | Use |
|---|---|---|
| GUS BC block F | GGCTACGGTCTCTCAAATGTTACGTCCTGTAGAAAC | Golden Gate |
| GUS BC block R | GGCTACGGTCTCCCGTATCATTGTTTGCCTCCCTGC | Golden Gate |
| PsNANA BC block F | GGCTACGGTCTCGCAAAATGATTTAGAGATGGGTT | Golden Gate |
| PsNANA BC block R | GGCTACGGTCTCGCGTATCAACATTTCTTAACCCTTC | Golden Gate |
| pEXPA AB block F | GGCTACGGTCTCTAAATCTGGTCTGTGAAAGAATTG | Golden Gate |

| | | |
|---|---|---|
| pEXPA AB block R | GGCTACGGTCTCTTTTGTTAAGATTCTGTATAAATTAATGAATG | Golden Gate |
| TCSn AB block F | GGCTACGGTCTCGAAATAAGCTTGACTAGTCAAAG | Golden Gate |
| TCSn AB block R | GGCTACGGTCTCCTTTGGGCAGGCCTGGATCCTCT | Golden Gate |
| PsNANA qPCR F | CCGAAATTCGGAAGCTTACTTTTA | qPCR |
| Ps NANA qPCR R | GATTTTTCCTAGCCTTGAGC | qPCR |
| PsTFIIa-F | CGGTGGAAATGCTGATGTTA | qPCR |
| PsTFIIa-R | GCTCCCTCCACATACCTCAA | qPCR |

**Table S2.** List of primers used for Golden Gate assembly and qPCR

**Methods S1.** Hairy root transformation with *Agrobacterium rhizogenes*. *Agrobacterium rhizogenes* strain ARqua1, containing the corresponding construct, was grown in lysogeny broth (LB) supplemented with tetracycline at a concentration of 10µg/mL for 24 hours at a temperature of 28°C. The bacterial culture was then collected and adjusted to an optical density (OD) of 0.4 in cold Murashige Skoog media.

For transformation, seeds were surface disinfected using 70% ethanol for 1 minute and then washed five times with sterile water and planted in a mixture of sterile vermiculite and gravel in a 1:1 ratio. This planting mixture was then topped up with autoclaved potting mix. Shoots from 14-day-old Torsdag plants and 17-day-old *na* plants were cut above node 2 in the case of Torsdag, and in the hypocotyl region for *na* plants.

The cut shoots were carefully placed in rock wool cubes that had been inoculated with 40mL of ARqua1 suspension. Excised shoots were maintained in a mini glasshouse with closed lid holes for 24 hours. After this initial period, the shoots were allowed to wilt without lids for another 24 hours. Following this, the plants were watered and covered once more for an additional 24 hours, all under controlled conditions of low temperature and light. The plants were then grown for a period of 10 days at a constant temperature of 21°C with 18 hours of light exposure. Following this growth period, the plants were transplanted into 1L pots containing a sterile mixture of vermiculite and gravel in a 1:1 ratio. During this transplant, any adventitious roots that had formed were dissected and removed, leaving only one root per plant to ensure optimal survival. After transplantation, the plants were inoculated with the RLV248G strain, and a nutrient solution containing 50mL of Long Ashton nutrient solution (Hewitt, 1966) with 2mM $KNO_3$, was applied to each pot. Long Ashton nutrient solution without nitrogen was provided on a weekly basis. The plants were harvested 5 weeks after transplantation for *NA* complementation experiments and 4 weeks post-transplantation for *TCSn::GUS* analysis.

**Methods S2.** Hormone analysis. For auxin and cytokinin extraction and quantification was carried out as outlined in (Bound et al., 2022). Stable isotope- labelled internal standards were [$^{13}$C6]-indole-3-acetic acid, and cytokinins: [$^{2}$H5] zeatin (Z), [$^{2}$H5] zeatin riboside (ZR), [$^{2}$H3] di hydrozeatin (DiHZ), [$^{2}$H3] di-hydrozeatin riboside (DiHZR), [$^{2}$H6] isopentenyl adenine (Isop adenine) and [$^{2}$H6] isopentenyl adenosine (Isop adenosine) (OlChemIm, Olomouc, Czech Republic) were added to each sample. For gibberellin quantification samples were extracted as outlined in (Ford et al., 2018). Internal standards ([$^{2}$H2]$GA_1$, [$^{2}$H2]$GA_8$, [$^{2}$H2]$GA_{12}$, [$^{2}$H2]$GA_{19}$, [$^{2}$H2]$GA_{20}$ and [$^{2}$H2]$GA_{53}$) were added to each sample. Samples were then analysed using a Waters Acquity H-Class UPLC instrument coupled to a Waters Xevo triple quadrupole mass spectrometer. For all samples, peak areas for endogenous and labelled hormones were compared and combined with the FW of samples to calculate ng/g FW.

**Methods S3.** Histochemical analysis for GUS expression. To stain for β-glucuronidase (GUS), roots where vacuum infiltrated for 20 minutes with staining solution ($NaPO_4$ 0.1M, EDTA 10mM, TritonX-100 0.1%, $K_3Fe(CN)_6$ 0.5Mm, $K_4Fe(CN)_6$ 0.5 mM and X-Gluc 2mM) and incubated for 6 to 10 hours at 37°C. After staining, roots were washed with phosphate buffered saline and stored in paraformaldehyde 1%. Section images were obtained by embedding the tissue in polyethylene glycol 1000 and sectioning 40µm slices on a microtome (Microm HM340E).
